# USE: An integrative suite for temporally-precise psychophysical experiments in virtual environments for human, nonhuman, and artificially intelligent agents

**DOI:** 10.1101/434944

**Authors:** Marcus. R. Watson, Voloh Benjamin, Thomas Christopher, Hasan Asif, Womelsdorf Thilo

**Affiliations:** Department of Biology, Centre for Vision Research, York University, Toronto, ON, M6J 1P3, Canada; Department of Psychology, Vanderbilt University, Nashville, TN, 37240; Department of Electrical Engineering and Computer Science, Vanderbilt University, Nashville, TN, 37240

**Keywords:** Psychophysics, Software, Cognition, Translational Neuroscience, Behavioral Control, Unity, Stimulus Presentation, Vision, Virtual Reality, Video Games, 3D, Virtual Environment, Experimental Design

## Abstract

1

**Background:** There is a growing interest in complex, active, and immersive behavioral neuroscience tasks. However, the development and control of such tasks present unique challenges.

**New Method:** The Unified Suite for Experiments (*USE*) is an integrated set of hardware and software tools for the design and control of behavioral neuroscience experiments. The software, developed using the Unity video game engine, supports both active tasks in immersive 3D environments and static 2D tasks used in more traditional visual experiments. The custom USE SyncBox hardware, based around an Arduino Mega2560 board, integrates and synchronizes multiple data streams from different pieces of experimental hardware. The suite addresses three key issues with developing cognitive neuroscience experiments in Unity: tight experimental control, accurate sub-ms timing, and accurate gaze target identification.

**Results:** USE is a flexible framework to realize experiments, enabling (i) nested control over complex tasks, (ii) flexible use of 3D or 2D scenes and objects, (iii) touchscreen-, button-, joystick- and gaze-based interaction, and (v) complete offline reconstruction of experiments for post-processing and temporal alignment of data streams.

**Comparison with Existing Methods:** Most existing experiment-creation tools are not designed to support the development of video-game-like tasks. Those that do use older or less popular video game engines as their base, and are not as feature-rich or enable as precise control over timing as USE.

**Conclusions:** USE provides an integrated, open source framework for a wide variety of active behavioral neuroscience experiments using human and nonhuman participants, and artificially-intelligent agents.

**Glossary:**

- **Active task**: Experimental tasks which involve some combination of realistic, usually moving, stimuli, continuous opportunities for action, ecologically valid tasks, complex behaviours, etc. Here, they are contrasted with **static tasks (see below)**
- **Arduino**: A multi-purpose generic micro-processor, here used to control inter-device communication and time synchronization.
- **Raycast**: A game-engine method that sends a vector between two points in a virtual three-dimensional environment, and returns the first object in that environment it hits. Often used to determine if a character in a game can see or shoot another character.
- **State Machine (also Finite State Machine)**: A way of conceptualizing and implementing control in software, such that at any one moment the software is in one, and only one, state. In **hierarchical state machines**, as used in the present software suite, these can be organized into different levels, such that each level can only be in one state, but a state can pass control to a lower level.
- **Static task**: Experimental tasks like those traditionally used in the cognitive neurosciences. Simple, usually stationary, stimuli, limited opportunities for action, simple behaviours, etc. Here, they are contrasted with **active tasks (see above)**.
- **Unity**: One of the most popular video game engines. Freely available.
- **Video game engine**: A software development kit designed to handle many of the common issues involved in creating video games, such as interfacing with controllers, simulating physical collisions and lighting, etc.

## 3 Introduction

### 3.1 Static and active tasks

Participants in most traditional psychology or neuroscience experiments are presented with impoverished stimuli and tightly-restricted response options. This approach, which for the sake of brevity we refer to as *static*, maximizes experimental control, constrains possible interpretations of results, allows for the comparatively easy creation of experimental tasks, and is a critical part of the reductionist approach that has led to many of the exceptional successes of the cognitive sciences. However, researchers are increasingly concerned with understanding behaviour and neural processing in tasks and contexts that are more complex and naturalistic, an approach we refer to as *active*. Here we present an integrated suite of software and hardware designed to aid in the development, control and analysis of active cognitive neuroscience experiments, the *Unified Suite for Experiments* (*USE*).

Given the success of static tasks and their dominance in the cognitive sciences, why should researchers bother with active tasks? We provide three answers to this question. First, there is the common concern about ecological validity and the generalizability of results, namely that results from simple static tasks do not necessarily generalize and may be misleading when taken to provide insight into real-world behaviour and neural activity (Kingstone, Smilek, & Eastwood, 2008; Schmuckler, 2001; Chaytor & Schmitter-Edgecombe, 2003). For example, there are substantial quantitative and qualitative differences in biases for gaze to be directed towards others’ eyes that depend on whether participants are viewing static pictures, watching movies, or are involved in genuine interactions with other humans (Risko, Laidlaw, Freeth, Foulsham, & Kingstone, 2012). By facilitating the generation of both static and active tasks using the same underlying task structure, USE enables the direct investigation of differences between them.

Second, more naturalistic stimuli and flexible possibilities of control can be more immersive and dramatically more motivating for human and nonhuman primate subjects (Bennett, Perkins, Tenpas, Reinebach, & Pierre, 2016; Bouchard, Guitard, Bernier, & Robillard, 2011; Slater & Wilbur, 1997; Witmer & Singer, 1998; Youngblut, 2007). As one testament to this, over the decades where video games developed more realistic visuals and more flexible and responsive controls, they developed from an obscure hobby to a multi-billion dollar industry with more consumers and larger revenues than film-making (The NDP Group, 2009; Shanley, 2017). As another testament, more realistic computer game environments can lead to reliable increases of learning outcomes in variety of contexts (e.g. Mayer, 2018).

Finally, active tasks enable the collection and exploration of a wide array of precise and multi-modal data about complex behaviors, which are necessary to generate hypotheses for understanding real-world behaviors (Kingstone et al., 2008). Despite more than two centuries of formal psychological research and millennia of informal speculation, there is no well-established body of fine-grained data about human action in most tasks. By providing such fine-grained data on realistic tasks, active tasks can produce data that enable both hypothesis-testing and exploratory or observational work, sometimes even in the same experiment.

### 3.2 Developing active tasks with USE

Active tasks may be appealing, but they are more challenging to develop, control, and analyze than static tasks. USE was developed to make these challenges more manageable. The practicality and scope of USE make it a versatile alternative to experimental creation and control suites such as the Psychtoolbox (Brainard, 1997; Kleiner, Brainard, & Pelli, 2007; Pelli, 1997), PsychoPy (Peirce, 2007; Peirce, 2008), or MonkeyLogic (Asaad & Eskandar, 2008a; Asaad & Eskandar, 2008b; Asaad, Santhanam, McClellan, & Freedman, 2013; Hwang, Mitz, & Murray, 2019), with the specific focus of creating, controlling and analyzing tasks that have the complexity, dynamism, and visual fidelity typical of video games. USE also provides unique solutions to common challenges of experimental control (see 5.2) that set it apart from other active experiment creation suites (Brookes et al., 2018; Doucet, Gulli, & Martinez-Trujillo, 2016; Jangraw, Johri, Gribetz, & Sajda, 2014).

USE provides an integrated solution for development, timekeeping, and analysis components required for typical experiments (**Figure 1**). The remainder of this paper delineates each of these components. For the development component, a set of scripts for the video game engine Unity enables the development of nested hierarchies of experimental control, as well as tools for common experimental requirements (data collection and recording, communication with other programs and equipment, etc). For the timekeeping component, the suite incorporates the *USE SyncBox*, an Arduino-based timing and I/O hardware system, to relay signals and codes between experimental hardware, and to maintain a central time record. This can send either event codes or simple pulses to other experimental equipment, and track the current monitor status using light sensors placed on the monitor, which allows the detection of any skipped or stuck frame (currently only in post-processing). Finally, for the analysis component, the suite includes a set of Matlab scripts for offline data parsing and timestream synchronization.

**Figure 1.**
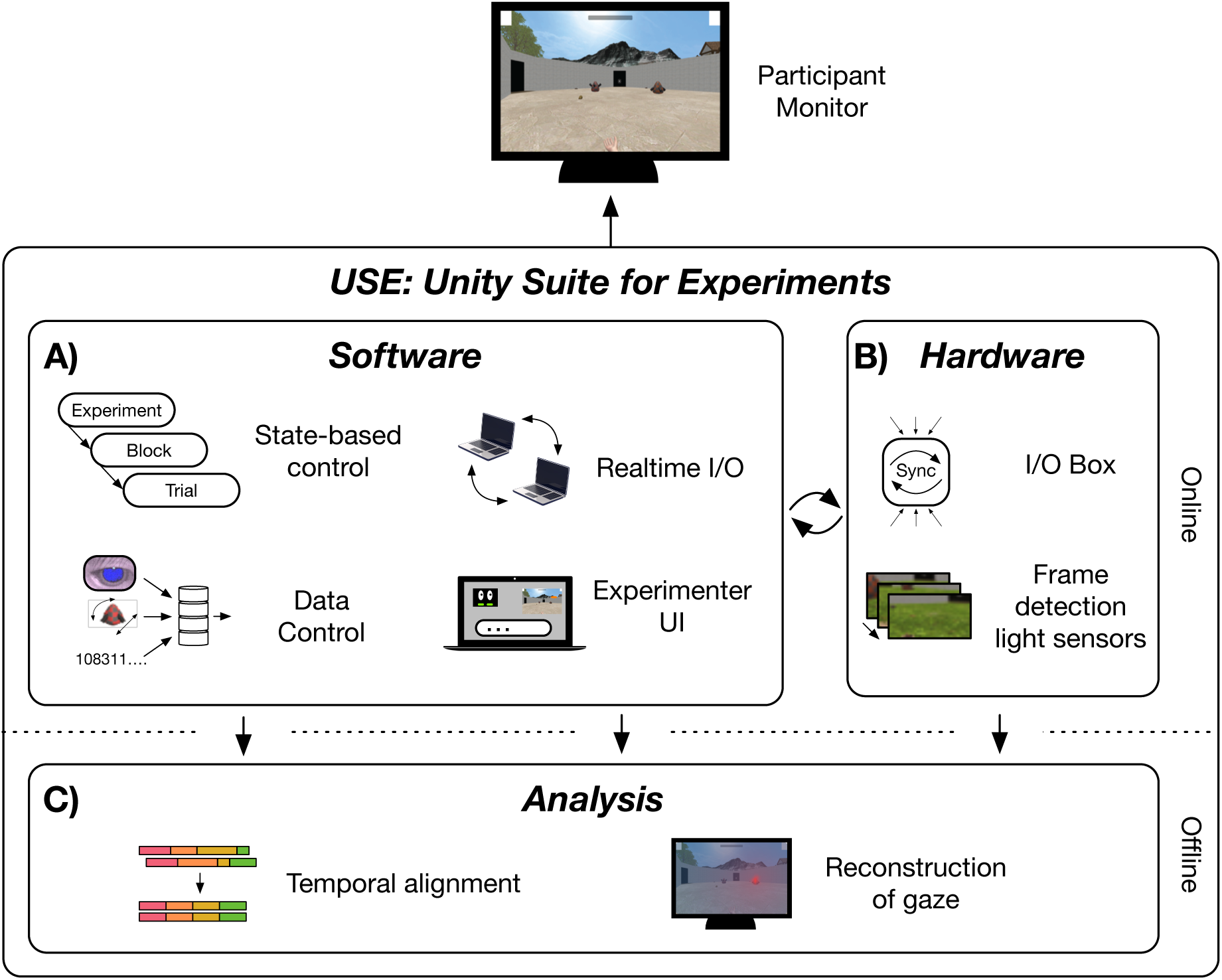
Unified Suite for Experiments allows strong experimental control in immersive tasks. Immersive, dynamic tasks present unique challenges to experimental control. To address these, USE includes novel contributions for (A) software, (B) Hardware, and (C) analysis tools. **(A)** USE employs a novel state-based control system that underpins its strong experimental control. Realtime I/O is based on current industry standards. USE also allows experimentalists to flexibly define and collate data occurring at multiple experimental timescales, and online visualization of subject’s performance. **(B)** The USE SyncBox is a newly developed, dedicated, and inexpensive machine that allows synchronization of different data streams (such as joystick data, eyetracker data, and game data), and can communicate with other hardware. As well, custom built photo-diode solution allows for the precise detection of physical frame onsets. **(C)** USE also contains a number of post-hoc analysis scripts, including a way to align all data sources, and reconstruct subject’s gaze.

USE allows flexible experimental protocols built around (i) active or static experiments, (ii) 2D and 3D displays, and (iii) touchscreen, joystick or keyboard/button press interfaces. In addition, it provides an interface for artificial agents that thereby can be tested with the identical settings used for experiments in humans or nonhuman primates. We anticipate that the key strength of USE is to facilitate the design and realization of active studies, though it can also be used to generate static tasks, as we have shown in prior studies (Oemisch, Watson, Womelsdorf, & Schubö, 2017; Watson, Voloh, Naghizadeh, & Womelsdorf, in press), enabling the comparison of active and static task variants.

Links to the key components of USE, as well as manuals, tutorials, and example experiments, can be found on our website (http://accl.psy.vanderbilt.edu/resources/analysis-tools/unifiedsuiteforexperiments).

### 3.3 Goals of USE

In developing USE, we aimed to satisfy multiple criteria. The goal was to develop a system that was:

- *Temporally accurate and precise* - all data should synchronize at millisecond precision.
- *Modular* - all software and hardware components are developed so that they can be used independently, or in different combinations.
- *Generic* - specific components can be easily adapted to multiple purposes.
- *Algorithmic* - each component of the system is explained in a principled way to facilitate implementation in other research contexts using other hardware (e.g. a different microcontroller board) or software (e.g. a different video game engine).
- *Translational* –functionally identical protocols can easily be generated for use with different groups of humans, non-humans, or artificially intelligent agents.
- *Offline reconstructable* - every monitor frame displayed during an experiment can be recreated at will in offline analysis and combined with synchronized information from any other data stream to reconstruct the experiment.
- *Cost efficient* - the software is all free, the total cost of the custom hardware is under $500, and experiments can be developed and run using any modern computer and consumer-grade monitor.
- *Multi-platform* - experiments can be developed on Mac OSX or Windows computers, and run on any modern computer (including Linux), smartphone, tablet, or gaming console, and can also be developed for the web.
- *Portable* - only a single computer, a small box for the arduino, two light sensors and a small number of cables are required for complete experimental control.
- *Practical* - the suite solves the key challenges in active experiment implementation and organization, and does so in ways that are intended to be as user-friendly as possible.
- *Open source* - all components of development and analysis software and I/O data streaming should be freely available under open source licenses.

One criteria we did *not* aim to satisfy was that developing experiments be a completely novice-friendly process. In our experience, suites which attempt to do so end up sacrificing generality, flexibility, and power. Thus, designing an experiment in USE, while it is much easier than doing so from scratch, does require a level of familiarity with both Unity and general programming principles. Actually running experiments created with USE, however, does not require any special knowledge and can be simple enough to be the responsibility of the undergraduates who are tasked with running more traditional studies in many laboratories (see 4.2.7). That said, we expect as more labs become involved in the development of active tasks, many programmatic components that define the actionable environment will be re-used as they are developed and shared through open-source collaboration, increasing the ease of experimental design.

### 3.4 The Unity game engine

The experiment development and control software components of USE are implemented in the Unity *game engine*. Game engines are development environments that implement various functionalities that games commonly require, such as the rendering of 2D and 3D graphics, physics simulation, realistic lighting, animation, sound management, etc. Unity is a free (but not open source) engine that runs on Windows, Macintosh, and Linux computers, supports building games for all major computers, phones, tablets, and game consoles, and has built-in support for stereoscopic presentation. Games made with Unity were downloaded over 5 billion times in Q3 2016 alone (Unity Technologies, 2017a). Some recent games of note made with Unity include Cites: Skylines (Colossal Order, 2015), Her Story (Barlow, 2015), Kerbal Space Program (Squad, 2015), Pokémon GO (Niantic, 2016), and Super Mario Run (Nintendo EDP, 2016). Aside from its price, Unity is attractive to small and medium-sized developers due to its relative ease of use, full feature set, comprehensive tutorials, and active online forums (Unity Technologies, 2017b). USE scripts are all written in C#, which as of 2019 is the only language supported by Unity.

### 3.5 Specific challenges of active tasks

Building an experimental suite on top of a game engine presents several unique challenges. First, game engines must ensure that all required commands are run in time for frame rendering to be carried out, but the precise execution order below the frame rate is often irrelevant, and commands can be spread across many scripts, making it difficult to follow their interactions. Command execution order in USE is guaranteed by a novel state-based system (cf. Wagner, Schmuki, Wagner & Wolstenholme, 2006) that enables the generation of nested hierarchies of control, such as the experiment-block-trial structure common in cognitive studies (section 4.2). As an example of the general utility of state-based control systems for experiment development, a state-based system has been used to create electrical engineering experiments in Unity exists (Liao & Qu, 2013), but its use case is sufficiently distinct from cognitive neuroscience that it is untransferable.

Second, just as control below the level of a frame can be difficult to achieve, accurate timing below the level of a frame is, for most practical purposes, unavailable within Unity. Indeed, it is difficult even to ascertain frame onset times, complicating synchronization of displays with other experimental hardware such as neural acquisition devices. To solve this, USE incorporates a newly designed, dedicated SyncBox hardware, which acts as a central timekeeper, a generic communication/synchronization device, and also uses data from light sensors to track the current frame status (section 4.3).

Finally, for studies that incorporate eyetracking, identifying gaze targets in a 3D scene rendered on a 2D screen is difficult, particularly when objects have complex shapes and the subject freely navigates through the 3D environment. In most static tasks, an Area of Interest (*AOI*) is defined around each object, with a larger radius than the object itself to account for measurement imprecision and the spatial extent of the fovea. This is much more difficult and computationally-intensive in active tasks, as objects’ two-dimensional silhouettes on the screen can be difficult to determine, and can change drastically from frame to frame. Our solution involves a novel method named *Shotgun Raycast*, which detects all objects whose two-dimensional silhouette falls within a specified number of degrees of visual angle from a gaze point on the screen (section 4.4).

These novel contributions extend Unity’s robust physics and rendering capabilities, and make it a suitable platform for behavioral neuroscience research.

### 3.6 Enabling translational research: human, non-human, and artificially intelligent agents

Our laboratory uses healthy human undergraduates, neurosurgery candidates, macaque monkeys, and artificial (reinforcement) learning agents to run tasks with a variety of different input mechanisms and recording devices. It was critical for us that USE be able to generate comparative task structure and data across all these groups. USE enables this in several ways. First, experimental protocols can be adjusted to meet the needs of different participant groups, such that different protocols using the same functional logic can be run from the same base scripts with small changes to configuration files (see 4.2.3). Second, an *input broker* enables entirely different inputs (for example, touchscreen presses, button clicks, fixations, or selection by an artificial agent) to produce the same effect in a trial (see 4.2.4). Finally, an artificial intelligence (AI) wrapper enables two-way communication with different learning agents (see 4.2.5), similar to existing AI testing platforms (e.g. Beattie et al., 2016; Leibo et al., 2018; OpenAI, 2018). If desired, full-resolution screenshots can be sent on every frame to visually-guided agents that can learn fully active tasks, as have proved influential in recent years (e.g. Mnih et al., 2015, Wang et al., 2018), as well as more traditional models that operate on simple digital vectors that represent static features of interest in the scene (e.g. Kruschke, 1992)

### 3.7 Online resources

Three Supplemental files are hosted with this paper:

- A review of the revised gaze classification algorithm used in our task (see 4.4.3 and 5.4).
- A detailed description of the timing tests performed on the USE SyncBox (see 5.1).
- A description of the normalization procedure used to produce Figure 6.

As previously noted, all USE manuals, scripts, schematics, and tutorial are available via the USE website (http://accl.psy.vanderbilt.edu/resources/analysis-tools/unifiedsuiteforexperiments/).

**Figure 2.**
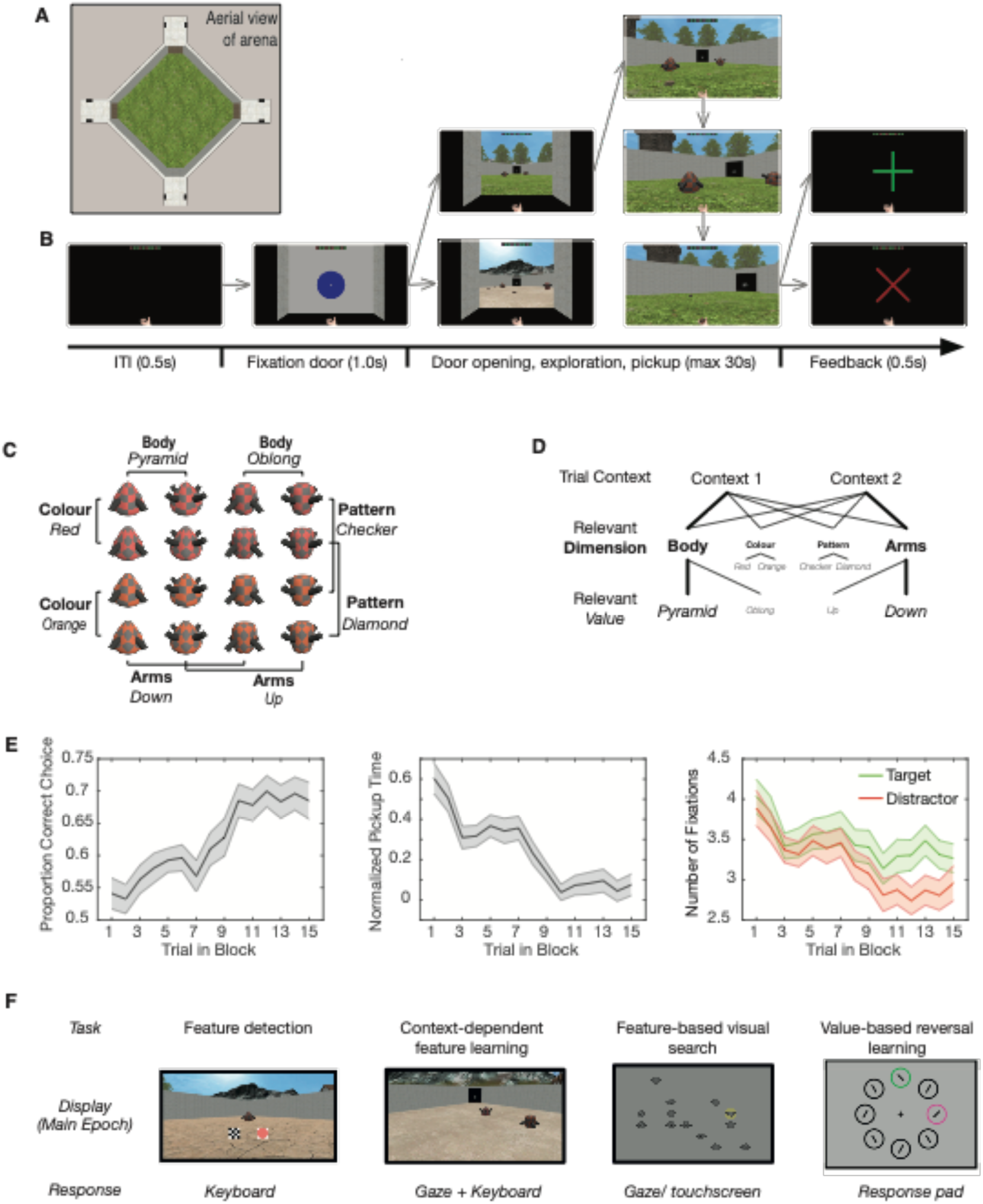
Example USE experiment. Details of one experiment coded in USE, and brief sketches of several others **(A)** An overhead view of the task arena. **(B)** An example trial sequence. Trials are separated by an ITI where most of the screen is black, but the avatar’s hand and their reward history is visible. Participants begin in one of the four corridors at the corners of the arena, facing a closed door. Fixating a white dot at the centre of a larger blue circle on the door opens it, revealing the arena. Participants then navigate towards one of the two objects in the arena and pick it up by walking over it. Finally, they take the object to any of the other corridor doors, where they are given visual feedback on their choice accuracy. **(C)** Set of objects used in the experiment (Watson et al 2018). Each object is defined by four feature dimensions (body shape, color, arm position, and pattern), each of which can take on two specific values (e.g. pyramidal or oblong, red or orange, etc). **(D)** An example reward structure that participants must learn in a given block. Objects are presented in one of two contexts, defined by the floor of the arena. Each context has an associated relevant feature dimension, and one rewarded feature value in that dimension. In this example, the objects presented in context 1 are rewarded if their body (feature dimension) is pyramidal (feature value). Thick lines and large font highlight the path towards the rewarded feature value, whereas small lines and font denote a path towards unrewarded values. **(E)** Accuracy, time from door opening to object pickup (normalized by dividing by the distance from door to object, and z-scoring within each participant), and number of fixations pre-pickup to rewarded target objects and unrewarded distractor objects. Shaded areas indicate standard error of the mean. **(F)** Other examples of experiments coded in USE. (Top) These tasks can have richly detailed or simple stimuli, in various degrees of interaction between player and environment. (Bottom) USE can support different inputs as required by the specific experiment.

**Figure 3.**
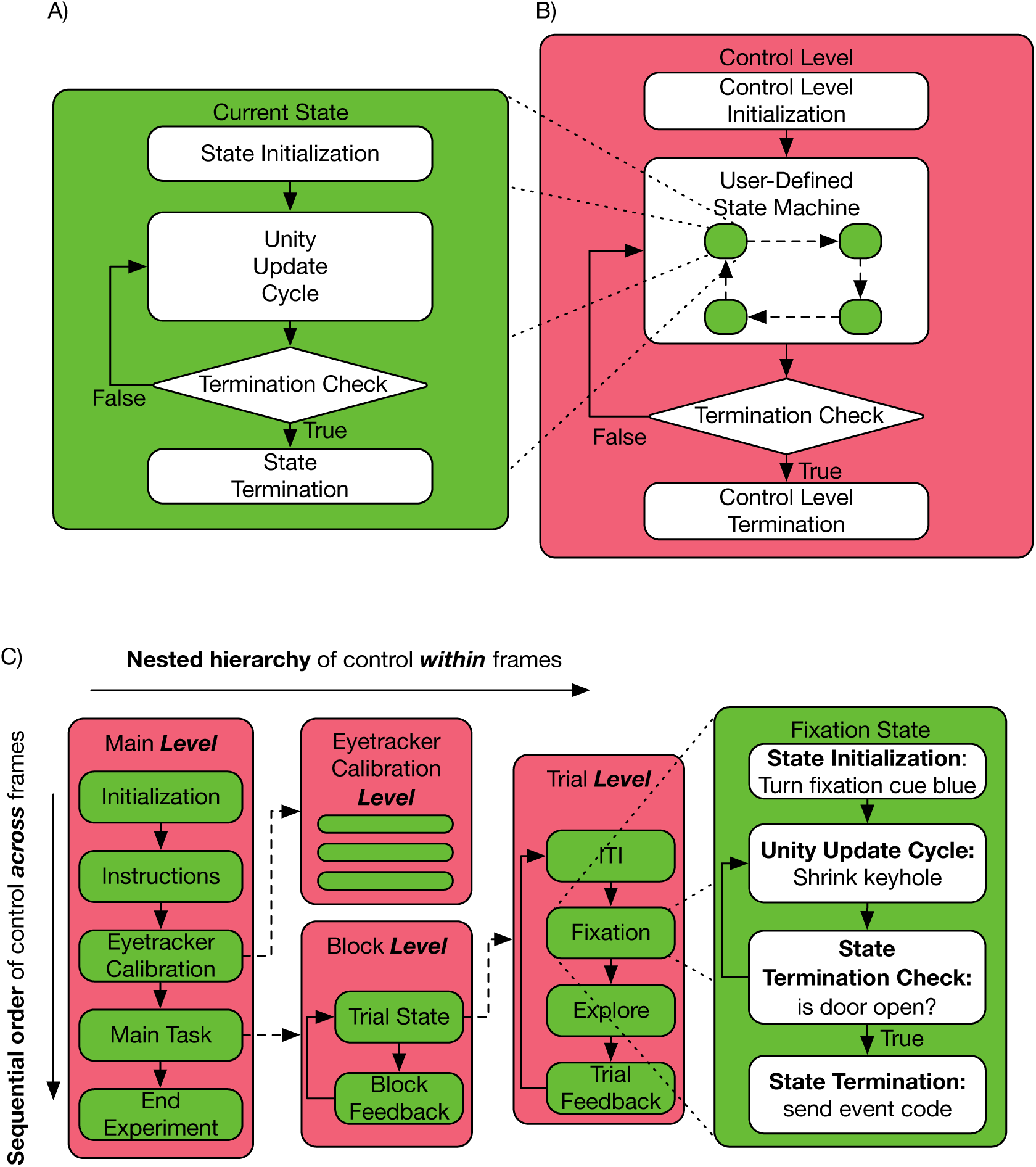
State/Level organization allows for flexible and strong experimental oversight. Depiction of (A) State and (B) Control Level architecture and their interaction in (C) an example experiment. White rounded rectangles denote groups of user-defined methods, diamonds represent Boolean conditions. Dashed lines with arrowheads represent the relationship of parent States and their child Control Levels. Dotted lines from all four corners of a State illustrate the zoomed-in view of the internal components of the state itself (not of child Control Levels). **(A)** A State contains an Initialization method group that runs only once at the start of the first frame in which the State is active. The Unity update cycle runs each frame until a Termination Check returns True, at which point the Termination method is run. (Actual States are somewhat more complex than illustrated, as it is possible to have an arbitrary number of Initializations, Termination Checks, and Terminations.) **(B)** A Control Level defines the States that can transition to each other. An Initialization method runs once the first time a Control Level becomes active. Only one state is active at any one time, and its Update/Termination Check cycle runs during each frame, after which the Control Level’s Termination Check is run, and if it returns true the Control Level’s Termination method runs. **(C)** Control flow of an example experiment, illustrating that Control Levels can be children of States. (This is only an illustrative portion of the full task hierarchy.) This allows nested control within frames, but also guarantees control across frames. The Main Level is the highest level of control. After completing an Initiation and Instruction State, the Level enters a Calibration State, which contains a lower-level Sequence that itself has multiple states. After completing this, the Main Task state is run, which also has a nested Block Level, which itself has a Trial State and associated Trial Sequence. Here, we explicitly illustrate the Fixation State within the Trial Sequence, and the processes operating during this State.

**Figure 4.**
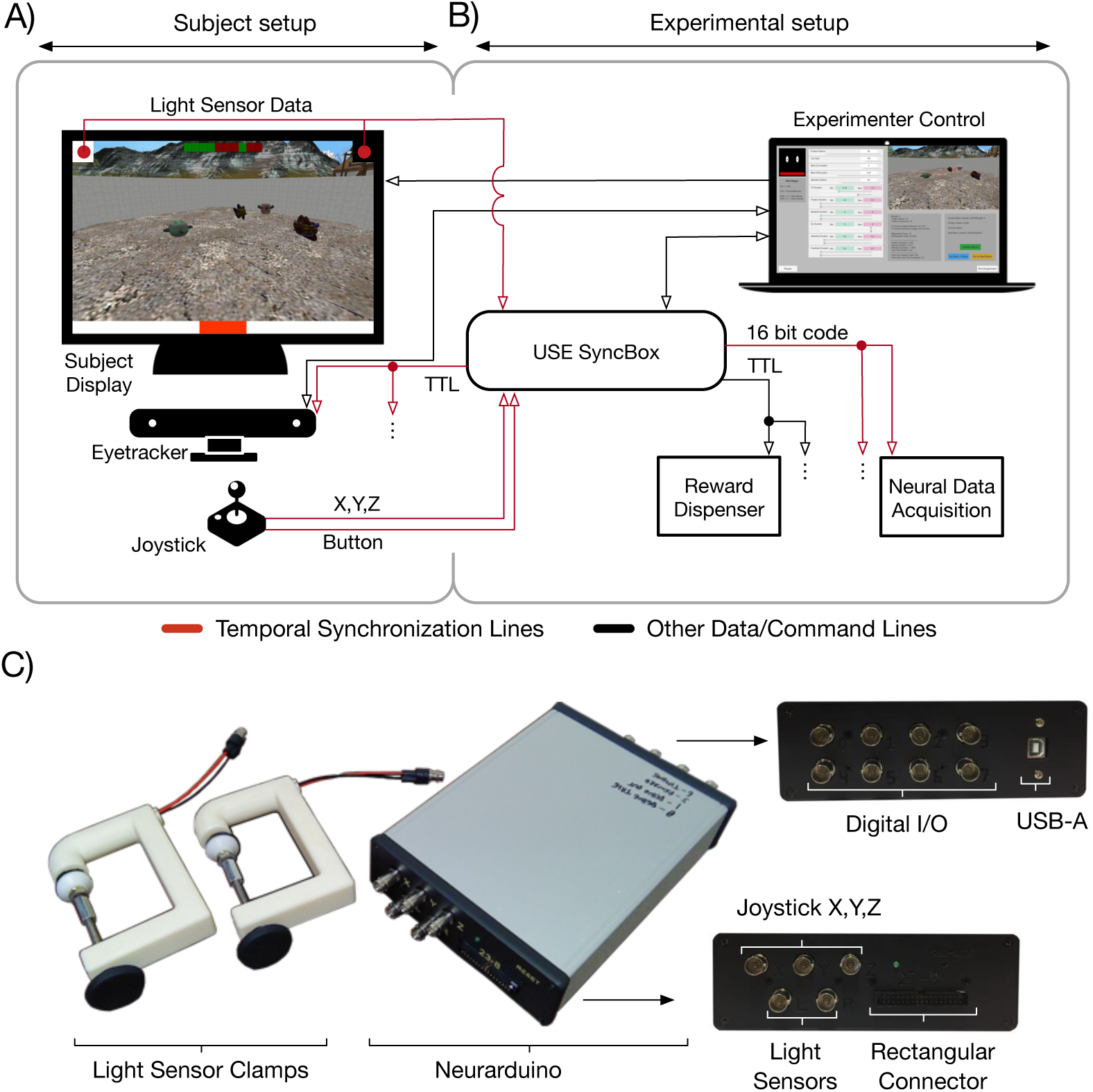
Example Connectivity diagram showing full capabilities of USE. A typical setup employing the full USE suite. **(A)** The participant setup (left) are those components of the experiment visible to participants, including the monitor, joystick, eyetracker, and light receptors. The virtual environment is displayed on the monitor, participants move through it using the joystick, and the eyetracker records their gaze behaviour. The monitor has patches in the top corners that change between black and white to indicate frame changes, which are picked up by the light receptors. The monitor is controlled directly by the experimental computer over its input line. Joystick output and photo-diode signals are communicated to the USE SyncBox. **(B)** The experimental setup includes the experimental computer, USE SyncBox, neural data acquisition device, and reward dispenser. The experimental computer runs the experiment, controls the monitor, and sends commands to the USE SyncBox. The USE SyncBox can forward event codes for time synchronization to the neural acquisition device, TTL pulses as needed to control a reward dispenser, or regular TTL pulses used for time synchronization to the eyetracker (TTL pulse). The experimental computer is directly connected to the participant display and eyetracker (1), all other communication is funneled through the USE SyncBox, whether from control computer to the peripheral device (2) or from the peripherals to the control computer (3). Lines denote communication from one component to another, with the arrowhead showing the direction of signals. Red is used to highlight those lines that allow post-hoc temporal alignment. **(C)** Photos of a USE SyncBox, its ports, and the custom clamp used to hold light receptors in place over participant monitors. All files and instructions needed to create these are available on the USE website.

**Figure 5.**
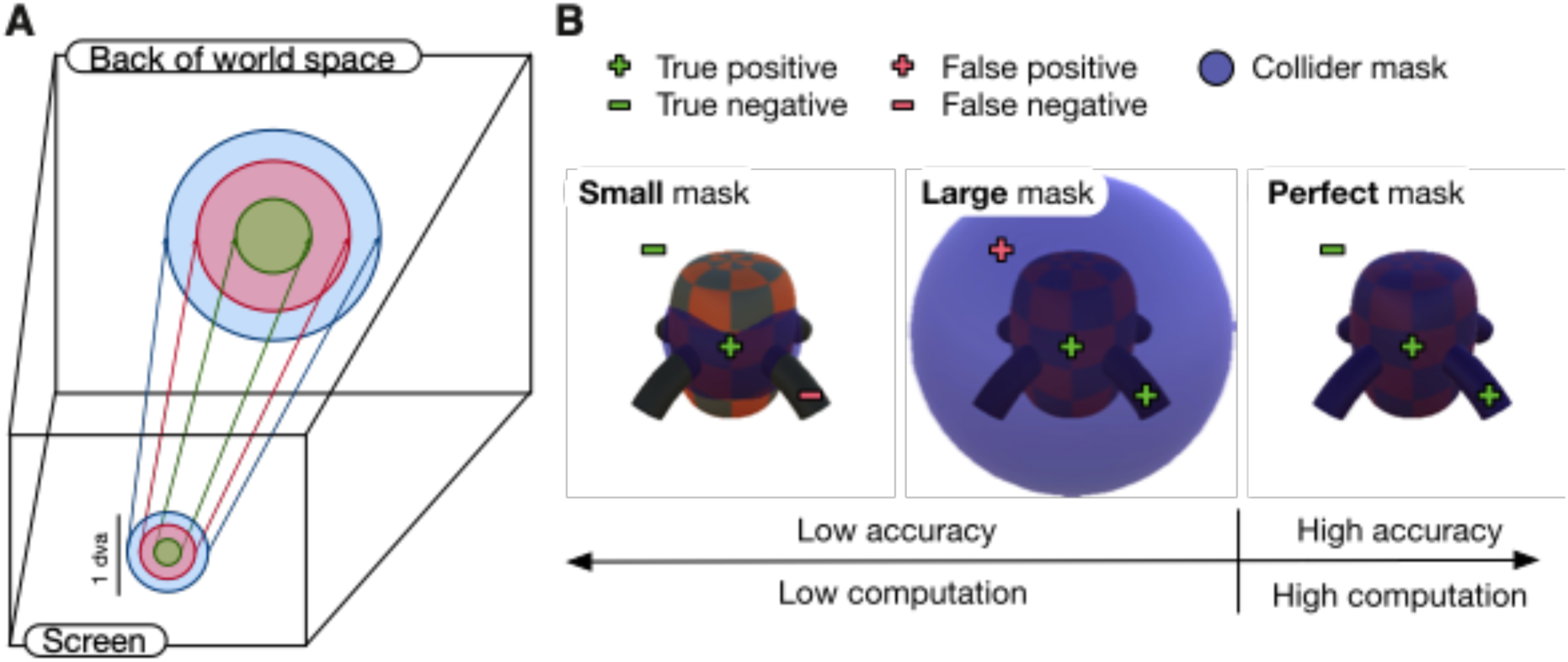
ShotgunRaycast allows gaze target determination in a 3D world. **(A)** Moving through a 3D world means that the size and shape of the silhouette of an object on the screen constantly changes, making it a challenge to determine gaze targets. The ShotgunRaycast solves this challenge. It defines a conic section, beginning with a circle on the camera and ending with a larger circle a long distance into the world space, which has the same center on the screen and whose projection onto the screen subtends exactly the same angle. This has the effect of finding any objects whose silhouette on the screen lies either completely or partially within the circle defining the smaller end of the conic section. Experimentalists define the density of sampling, and radius of the circle in degrees of visual angle. **(B)** Detection of an object depends on the shape of the collider associated with it. Simple colliders (e.g. spheres; left, middle) are computationally inexpensive. However, they are inaccurate, either because raycasts miss the object - resulting in false negatives (left) - or raycasts hit the outside of an object - resulting in false positives (middle). Alternatively, mesh colliders that perfectly match the shape of an object are perfectly accurate, but are computationally expensive when the object is defined by a high number of polygons.

**Figure 6.**
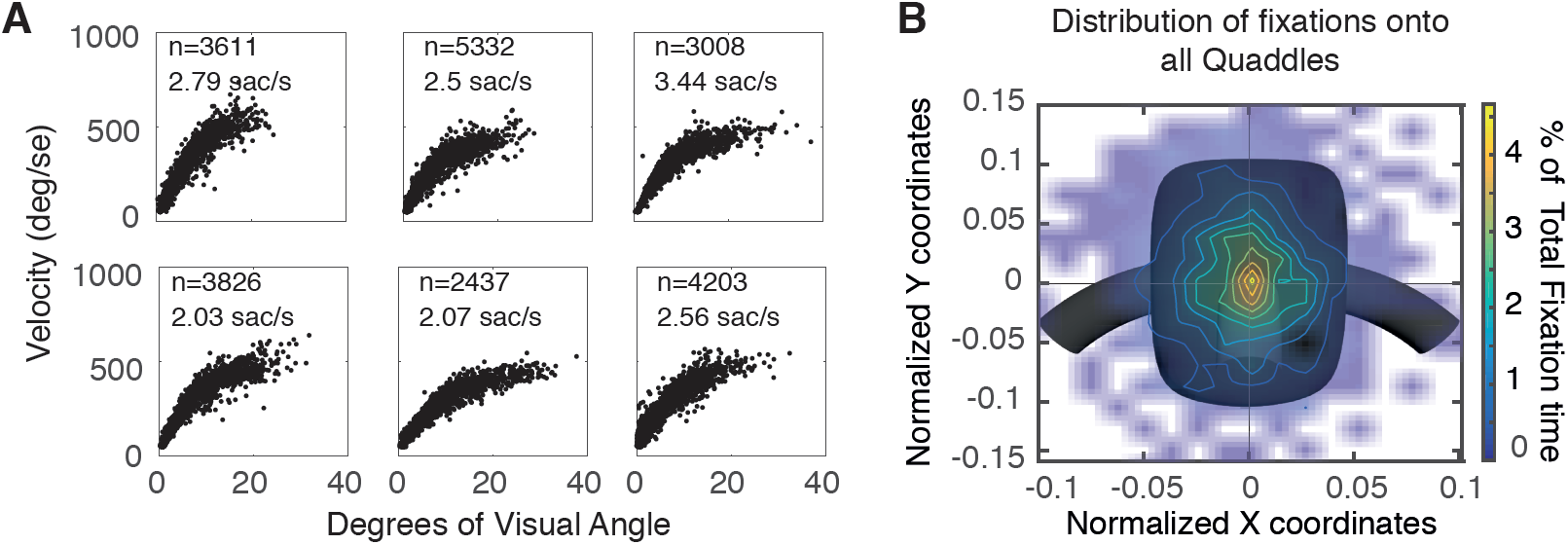
Gaze behavior during example experiment. **(A)** Saccadic main sequence of six representative participants performing the context-dependent feature learning task. Saccade rate ranged from ∼2-3.5/s. An inflection point in the main sequence occurs around 10-15 deg. **(B)** Density of fixations onto a standardized object for one participant (see methods). Fixation centers were normalized for change in object size with depth and mapped onto a standard object located 4 world units away from the camera. Details of the normalization process are in Supplemental Materials 3. Color indicates the relative amount of time that participants spent fixating each point. Participants were most likely to fixate close to object centers.

### 3.8 Outline

In the remainder of the paper we outline an experiment that provides specific examples of the various components of USE in action (4.1), before turning to an overview of USE’s experimental design and control software (4.2), timing and communication hardware (4.3), and offline analysis capabilities (4.4). We then present results demonstrating the robust timing and control capabilities of USE (5) and discuss the implications of this work (6).

## 4 Methods

### 4.1 Example experiment

Static tasks built with USE have already been published (Oemisch et al., 2017; Watson et al., *in press*), but to demonstrate more of the suite’s capabilities, we briefly present an active learning task, in which participants must navigate through a 3D scene, interact with objects, and try to learn a reward structure. A full overview of the task and results will be presented in another manuscript (Watson, Voloh, Naghizadeh, & Womelsdorf, *in preparation*), here we focus on reporting sufficient detail to show USE’s capabilities.

Figures 2a and 2b display the task environment and trial protocol. In brief, participants must choose between two objects on each trial, one of which is rewarded and one of which is not, and they must learn the rules that govern reward through trial-and-error. Trials consist of participants navigating through an arena using a joystick, picking up one of the two objects by walking over it, and taking it to a door where they receive visual feedback on choice accuracy. The two objects are selected from a pool of 16 (Figure 2c, cf. Watson et al., *in press*), which are parametrically-defined by four feature dimensions (shape, colour, pattern, and arm direction) of two possible values each. In each block there are two rules, both of which reward a single feature value, but each of which only applies in a single context (symbolized by the arena floor). For example, objects with pyramidal body shape might be rewarded on grass floors, while objects with downward-bending arms rewarded on marble floors (Figure 2d). Once participants have achieved satisfactory performance on a block, new rules are generated, and a new block begins. Participants are explicitly informed of each block change, and told that they will now have to discover a new pair of rules.

50 participants ran in the study for approximately one hour each. Two participants were excluded due to chance performance. Gaze was tracked using a Tobii TX300 combined head-free eyetracker and OLED monitor (Tobii Technology, Inc.). Figure 2e demonstrates that over the course of a block they learned to make more accurate choices, to make these choices more rapidly, and to preferentially fixate the rewarded object over the unrewarded one (see 4.4.3 for details of how fixation targets are determined in USE).

Figure 2F surveys the display configuration, structure, and response mode of several of the other experiments that have already been coded in USE.

### 4.2 USE: Unity Suite for Experiments

#### 4.2.1 States and Control Levels

Most software commands in USE are controlled by an architecture of *States* and *Control Levels*, which allow experiments to be defined as a series of *hierarchical finite state machines* (Wagner et al, 2006). This State-Level architecture is flexible enough to support any standard experimental hierarchies such as experiment-block-trial, as well as more complex structures. Here, we explain their abstract functional role and give examples of their use in the experiment described in 4.1.

A State is an object (in the object-oriented programming sense) that defines the operation of the experiment during a period of time (Figure 3a). These are organized in Control Levels that group States that operate at similar levels of abstraction in the experiment (Figure 3b). In the example experiment, a Trial Level groups together the States that define the various trial epochs, while a Block Level groups together States that define the block sequence, and passes control to the lower Trial Level as needed, and an even higher Main Level groups the States that govern aspects of the experiment outside the Block (Figure 3c). Thus, States can be used to define both the finest-grained and the most general parts of the experiment.

Each State specifies commands that are run every frame while it is active, and initialization and termination commands to be run at the start of its first and the end of its last frame, respectively. For example, the Fixation State of the example experiment’s Trial Level determines what happens when the participant stands in front of a door before entering the arena. At the start of the first frame in which an experiment enters a new State, a `StateInitialization` method runs. For the Fixation State, this method turns a circle on the door blue, cueing the participant to look at it. Within each frame of a State, a number of method groups run, controlling Unity’s update cycle (for full details of the update cycle, see the USE manual). For the Fixation State, these methods control the size of the blue circle, which shrinks as it is fixated, and the position of the door, which opens after the circle has completely shrunk. The update cycle is followed by `StateTerminationCheck` methods that verify whether any end conditions for the state have been reached. For the Fixation State, there is a single condition: is the door fully open? If one of the end conditions has been met, a `StateTermination` method group is run at the end of the last frame of the state, and the experiment transitions to a new State on the following frame. If no end conditions have been met, the current State’s update cycle methods run again on the following frame. For the Fixation State, the `StateTermination` method starts opening the door, revealing the main arena behind it, and on the following frame the Trial Level transitions to the Explore State.

States are grouped together in Control Levels (Figure 3b). Any State in a Control Level can transition to any other State in the same Level. These transitions are defined by `TerminationSpecifications`, each of which includes (a) a `StateTerminationCheck`, (b) a `StateTermination`, (c) a successor State, and (d) the `StateInitialization` of the successor State that will be run. By defining the States that make up a Level, and the desired transitions between them, an experimenter has defined the finite state machine that constitutes that Level.

Like States, Control Levels have Initialization and Termination methods. `LevelInitialization` and `LevelTermination` methods run once each, at the start of the first frame (prior to any `StateInitializations`) and end of the last frame (after any `StateTerminations`) in the Control Level, respectively.

#### 4.2.2 Hierarchical State/Level organization

A critical aspect of USE is that *Control Levels can be children of States*, allowing Control Levels to pass control to sub-Levels. This guarantees that commands run within the experimenter-defined order, ensuring that the subject’s experience is as the experimenter had intended.

The State/Level architecture governing the example experiment (4.1) is visualized in Figure 3c. The Main Level is the top level of the hierarchy, which defines high-level States such as setup, eyetracker (gaze) calibration, and the task itself. The eye tracker State has an associated Eyetracker calibration Level, which itself has multiple States associated with it (presentation of the calibration dot, analysis of the estimated gaze location, experimenter acceptance or rejection of the calibration results, etc). Once this has been completed, we progress to the Main Task state. This has embedded within it a Block Level, which itself has a Trial State that passes control to a Trial Level. In this example, there are four trial States – inter-trial interval, fixation, exploration, and feedback — each of which will include its own initialization, update and termination methods. We have described these in detail for the Fixation State above.

Once all trials in a block have been completed, the Block Level’s `TerminationCheck` verifies if there are blocks remaining in the experiment. If so, the Block Level loops back to the beginning, and a new round of trials begin, but if not, the Block Level ends and the Main Level shifts to an End Experiment State, which performs various housekeeping functions before closing the application.

We anticipate that the large majority of experiments will use at least a two-level hierarchy of Block and Trial Level, and that more complex experiments will benefit from a top-level Main Level, as used in the example experiment. State and Level logic provides a flexible way of designing experimental hierarchies, while maintaining precise control over the order of commands during Unity’s primary update loops.

#### 4.2.3 Selective reuse of States for different experimental protocols

It is quite common, particularly in translational research, for modifications of the same basic experimental protocol to be used in different experiments. For example, non-human primates typically require a fluid or pellet reward in addition to visual feedback for correct performance, while artificial agents might require only an abstract numeric representation of stimuli, without any visual input at all. Such modifications are often implemented by copying and pasting large chunks of code between experiments, which inevitably results in unintended differences, as later changes do not propagate across the different versions. Another solution is to gate portions of code using *if* or *switch* statements, which quickly makes code extremely hard to follow as the number of cases mounts.

USE simplifies this process by enabling experimenters to define more States than are necessary for a single experiment, and to select which of these States will be included in a given Control Level at runtime simply by listing their names when defining States. Thus, it is easy to generate highly similar experiments that differ only in a few trial or block States, guaranteeing that the logic they share will remain constant even after later edits to code.

#### 4.2.4 Input broker

Changes to experimental protocols often require very different actions to have similar effects. For example, our laboratory has run studies where objects are selected by keyboard button presses, touchscreen touches, fixations, or with the output of an artificial learning agent (*see* Figure 2f). A dedicated input broker is used to handle all of these cases and can be customized to support whatever input methods are needed for a given experiment.

#### 4.2.5 AI wrapper

Artificial learning agents implemented in any language can interact with USE and play through the same experiments as human or animal participants, using the suite’s AI wrapper. These agents can act based on representations of the environment that consist either of a numerical vector (e.g. different object feature values would be encoded as different numbers), or a screenshot of the current frame. To integrate an AI with a USE task, three core functions must be implemented: 1) `Reset` – starts/restarts the task, and configures the wrapper to use numerical representation or screenshots to represent the environment. 2) `Step` - moves the environment to the next step, and outputs its numerical or image representation. 3) `Act` - takes an action as an input parameter and returns the result of applying this action on the environment, including reward value and indicators of whether the current trial, block, or experiment has ended.

The USE suite includes a Unity-hosted TCP server to serve AI player requests, and a python library that implements a TCP client. Any python-based artificial learning agent that can operate using the Step and Act functions described above can interface with the client, and thus run any tasks developed in USE. The tutorial includes a simple python agent as a demonstration of these capabilities.

#### 4.2.6 Data control

USE incorporates a generic `DataController` class that enables the flexible collection and writing of as many data streams as may be desired, and writes these to text files. These can include variables of any type, and each `DataController` object is independently controlled, meaning that data can be collected and written at different frequencies. For example, it makes sense to collect positions of the camera and moveable objects every frame, while trial accuracy might only be updated once every few seconds, and block-level information might be generated every few minutes.

Importantly for timing and later analysis purposes, in our studies we generate FrameData files, which are updated every frame and include the positions, rotations and sizes of all objects for which these values can change (see 4.4.2), as well as the current expected state for each of two flickering patches beneath the light sensors used for timing alignment (see 4.4.1).

#### 4.2.7 Initialization Screen and Experimenter View

When running a USE experiment, our laboratory employs a two-monitor setup, one for the experimenter and one for the participant (see Figure 4). The experimenter’s screen allows various factors to be specified both at the start of an experiment and during its runtime. These are intended to allow individuals who may not have the programming experience to develop a task to nevertheless be able to control it at runtime, as needed in many laboratories.

At the start of an experiment, a customizable initialization screen is displayed. This can include file selection dialog panels enabling the selection of configuration files, as well as text boxes, Boolean check boxes, and numeric sliders, to specify information that might be desired for the experiment (e.g. subject ID, condition, or duration of different trial States). Each of these can be set to display the previously-chosen value as a default, or some function on this (e.g. subject numbers can be set to automatically iterate with each session).

Throughout an experimental session, the experimenter’s monitor displays three main components: (1) a panel showing a stream of exactly what the participant is viewing on their monitor, with overlaid real-time gaze or touch positions if desired, (2) text panels displaying real-time information of various kinds, such as messages from hardware, or summary information about participant performance, and (3) text dialog boxes, Boolean check boxes, and sliders. The interactive components in (3) can be used to modify experimental variables in real-time.

All components of both the initialization and experimenter view can be customized using Unity’s editor and external configuration files.

#### 4.2.8 Realtime I/O

USE handles real-time communications with other software and hardware via `SerialPortController` and `UDPPortController` objects, which include methods for setting up ports, reading incoming data from system buffers, storing it in USE-specific buffers for use by other methods, and clearing these buffers. For instance, we communicate with a python script controlling our eyetracker via UDP, and with the SyncBox via serial. One special case of serial data are event codes, which are handled by a specific `EventCodeController` and sent to the SyncBox. Typically, codes are prepared a frame in advance, and sent as quickly as possible after the new frame onset. However, Unity’s internal limitations mean that the latency between frame onset and event codes is less stable than might be desired, and for millisecond precision event code timestamps are adjusted with automatized scripts in offline post-processing (see 5.3).

### 4.3 The USE SyncBox – timekeeping and communication

The USE SyncBox is used to channel communication between experimental hardware during an experimental session, and to generate a single, highly accurate timing record that enables the time-alignment of the various data streams produced by these different pieces of hardware, as well as the physical onset of frames on an experimental display. It consists of a commercial Arduino Mega2560 r3 board with custom firmware and a custom shield connecting it to a number of I/O ports. Figure 4 shows how the box might be used in a typical experiment.

In the laboratories we have used the SyncBox, it has been used to send event codes to different neural acquisition devices (Neuralynx, BrainAmp, and Neuroscan systems), simple timing pulses to Tobii eyetrackers, and commands to control fluid and pellet dispensers used to reward non-human primates for performance on tasks. It has also been used to receive signals from the light sensors described in 4.3.1, as well as from a custom joystick. Other I/O capabilities can be added by modifying the firmware, and, if necessary, building an adapter to modify the layout of the event code lines. The SyncBox is thus a multi-purpose I/O device that can be used in a variety of experimental settings and quickly adapted between setups.

The two-way connection to the experimental computer is over a USB serial port. There are eight digital I/O channels connected to BNC jacks, five analog input channels also connected to BNC jacks, and 16 digital output channels connected to a 34-pin rectangular connector. Two of the analog inputs are dedicated to receiving data from light sensors attached to the participant monitor, which are fed through a pre-amplifier circuit which performs amplification, low-pass filtering, and DC subtraction on the light sensor signals (see the Syncbox Manual). In the example experiment (4.1), one digital channel was used to output a timing pulse every second that was received by a Tobii TX300 eyetracker, while the rectangular port was used to send event codes to a Neuralynx Digital SX Lynx SDSata acquisition system. Full details of all hardware are available on the USE website.

Full firmware details and code are also available on the USE website. In brief, the box runs an *interrupt-driven loop* every 0.1 ms, and a *host communication loop* that runs as quickly as possible, but without guaranteed timing. The host loop controls all communication over the serial port with the experimental computer, The interrupt-driven loop reads from and writes to all other inputs/outputs and maintains information about scheduled events.

The host computer can issue commands to the box over the serial port to control most aspects of its functioning, including event timing and the interval at which data is reported. The example experiment (4.1) received data with a sampling interval of 3.3 ms, and faster reports are possible. Details of the box’s timing capabilities are found in 5.1.

#### 4.3.1 Frame detection using light sensors

To precisely track frame onsets and detect skipped frames (where an expected frame is not rendered) or stuck frames (where a single frame persists for two or more frames), we place the box’s light sensors over two small patches at the top corners of the experimental display (see Figure 4). One patch changes from black to white every frame (*timing patch*), while the other encodes a binary sequence (*coding patch*), specifically the 24-character sequence consisting of the 3-digit binary representations of the numbers 0-7. Deviations from the expected timing and sequence of blacks and whites can then be detected, and skipped or stuck frames identified (5.2). Light sensors were housed in custom 3D-printed clamps that fit a wide range of monitors (.STL files for 3D printing available on our website).

#### 4.3.2 Time synchronization with external devices

There are three ways of syncing the time streams of external devices with the USE SyncBox, and thereby the rest of the experimental equipment. First, devices can receive up to 16-bit event codes sent on command over the USE SyncBox’s rectangular port (e.g. the Neural Data Acquisition box in Figure 4). Second, they could receive regular pulses sent over one of the USE SyncBox’s single digital ports (e.g. the eyetracker in Figure 4). Finally, they could themselves send data to the USE SyncBox (e.g. the joystick in Figure 4). Custom adapters may need to be built for any of these purposes, and we provide schematics of adapters that alter the rectangular port’s output for Neuroscan and BrainAmp acquisition systems on the USE website. Connecting an external device’s output to the USE SyncBox may also require modifying the USE SyncBox’s firmware.

If devices are not connected to the USE SyncBox in this manner, their time synchronization will be limited by Unity’s update cycle. Thus, a standard consumer-grade joystick or keyboard connected over USB will have an unavoidable jitter of up to a frame (16.7 ms on a standard monitor), as well as any delays introduced by the ports themselves.

### 4.4 Analysis Pipeline

#### 4.4.1 Time-alignment of data files

The various data files produced during an experiment are produced by devices using different clocks, which all need to be aligned to a single timestream. In the current USE version, this happens offline. Generally speaking, experimental hardware receives timing pulses or event codes from, or sends digital signals to, the USE SyncBox. Alignment is then a simple matter of assigning USE SyncBox timestamps to each matching code or pulse, either received or generated, in the other hardware’s data.

Alignment with data produced internally by the host computer, and with the physical state of monitor frames, is performed in a separate script. The frame-by-frame data stored during the experiment (3.2.7) includes the putative current status (black or white) of each of the two patches under the light sensors connected to the USE SyncBox, as reported by Unity. By analyzing the sensor voltages over the course of the experiment, the physical state of these patches on each frame is determined (see 5.2). Any skipped or stuck frames can then be identified by comparing the states of the clock and signal sensor, and the physical onsets of each frame, as opposed to Unity’s estimated onset times, are determined. The frame data can then be modified to reflect the actual status of every single frame in the experiment. At this point, time alignment of all data is complete, as all datastreams are referenced to the USE SyncBox’s timestamps, and can thus be directly compared with each other.

#### 4.4.2 Experimental session reconstruction

During a typical experimental session, the frame-by-frame position, rotation, scale, and other properties of interest for each object in the scene is recorded in a *FrameData* file (4.2.6). This enables the complete reconstruction of the experimental session. During reconstruction, Unity’s physics engine is ignored, and instead each object is directly assigned its properties as recorded. Each replayed frame contains a perfect re-presentation of the three-dimensional scene, including the camera’s position, and thus the image on the monitor is identical to that originally seen by the participant. A video of the entire experimental session, or simply particular moments of interest, can be generated offline, with no need to record during the session itself. In the current version of the suite, this frame-by-frame reconstruction requires custom code modifications for each experiment, a process that we intend to streamline in future version updates.

Other suites have replay capabilities (e.g. Jangraw et al, 2014), which USE extends in several ways. First, gaze positions, mouse or touchscreen clicks, or data recorded by other equipment during the session can be overlaid on the screen. Also, since all skipped and stuck frames are known (see 4.3.1), replays are perfect representations of what was actually displayed. Individual frames can be exported at full or reduced size and analyzed at will, for example using saliency estimation algorithms (e.g. Borji, Sihite, & Itti, 2014). Finally, gaze targets can be determined, as described in the following section.

#### 4.4.3 Gaze target determination

We smooth and classify gaze data using a modified version of an existing algorithm that provides superior processing of active gaze data (Andersson, Larsson, Holmqvist, Stridh, & Nyström, 2016; Corrigan, Gulli, Doucet, & Martinez-Trujillo, 2017; Larsson, Nyström, & Stridh, 2013; Larsson, Nyström, & Stridh, 2014; Nyström & Holmqvist, 2010), adapted to handle noisier data by using estimated angular acceleration (Engbert & Kliegl, 2003; Engbert & Mergenthaler, 2006) and robust statistics (Leys et al. 2013; Wilcox, 2010). Supplementary Methods 1 provides details of the algorithm, which is described further in (Voloh et al., submitted). The results of this classification are detailed in 5.4 below.

After smoothing and classification, gaze targets can be determined (Figure 5). The logic here is something like the inverse of gaze target determination in typical static tasks, in which an AOI is specified around an object of interest, and gaze points that land within this AOI are treated as landing on the object. In USE, any object whose two-dimensional silhouette on the screen lies within a specified distance from each gaze point is treated as a potential target of that point. For any frame of interest, the corresponding gaze points are identified, and a `ShotgunRaycast` method is used to determine which objects have been foveated (Figure 5A). This defines a conical section filled with multiple raycasts in Unity’s three-dimensional worldspace, whose smaller end is a circle centered on the gaze point, placed exactly on the camera (i.e. the surface of the screen). The larger end is defined such that its silhouette on the camera is identical in size and location to the smaller end. Thus, if any of the rays intersect with an object, its two-dimensional silhouette on the screen lies within the circle defined by the smaller end, i.e. is within the desired degrees of visual angle of the gaze point. The function returns a complete list of all objects hit by the rays making up the conical section. The density of sampling and the radius of the circle surrounding the gaze point are experimenter-defined.

Whether a gaze target is reliably detected depends on the shape of its associated mask, or *collider*, which defines the surfaces that raycasts can hit (Figure 5B). A mesh collider that perfectly matches the shape of the object will also result in perfect detection, but for shapes defined by a high number of polygons, mesh colliders are computationally expensive. Simpler shapes (usually spheres) are computationally cheap but involve a tradeoff in detection accuracy. Various intermediate collider types can be used to define shapes with greater or lesser degrees of fidelity to the precise object shape. Which one experimenters use will depend on their particular needs. For most of the 3D objects used in our studies (Figure 2C; Watson et al, in press), the high fidelity afforded by a mesh collider does not come at a high enough computational cost to warrant concern.

After replaying the session using shotgun raycasts for each gaze point, the experimenter then has the sample-by-sample specification of all objects falling within the desired degrees of visual angle from each gaze point, which can be further analyzed according to gaze type (e.g. fixations, smooth pursuits), or objects, as required by the experiment (see 5.4).

## 5 Results

### 5.1 USE SyncBox timing

The USE SyncBox enables digital signals to be generated with low latency and high precision (on the order of 0.1 ms for both). Specifically, digital timing pulses are generated with a clock stability of 100 parts per million and jitter between pulses of approximately +/- 0.02 ms. When commanded to send a single digital pulse or event code, the delay between trigger and output is 0.01 to 0.11 ms, limited by the box’s scheduling interval of 0.1 ms. Analog signals are digitized at 1 ms intervals with sample timing known to 0.1 ms accuracy. Light sensor signals pass through a pre-amplifier before digitization, which causes an additional delay of 0.33 ms.

Full details of the tests supporting these timing specifications are found in the Supplementary Material 2.

### 5.2 Frame detection

To quantify the performance of frame detection and frame onset determination for a monitor with 60 Hz frame rate, signals were recorded from the clock and signal light sensors over approximately 1.5 hours. The data for each sensor was analyzed by identifying peaks and troughs in the first derivative of the voltage trace, where intervals peak-to-trough were classified as black frames, while trough-to-peaks were classified as white. The median duration of black or white periods in the clock signal was 16.7 ms. In the entire test period, there were exactly 32 periods (of a total of 336,722, ∼0.0001%) whose duration was not within 3.3 ms (one reporting interval) of 16.7, 33.3, 50.0, or 66.7 ms. Thus, effectively all white and black periods have a duration that corresponds to an integer multiple of the expected duration of a single frame on a 60 Hz monitor. Any frame where this multiple is greater than 1 constitutes a skipped or a stuck frame, which can be identified by investigating the corresponding *coding patch’s* data.

### 5.3 Frame onset to event code timing

A critical question for many neuroscientists will be *what is the relationship between the physical onset of a frame and timing signals sent on that frame?* We tested this by connecting an oscilloscope to (a) light sensor placed on the host computer’s monitor, and (b) one line of the USE SyncBox’s rectangular port. A USE script was written to alternate the patch of the monitor below the light sensor between black and white every frame, and to send an event code as rapidly as possible at the start of each frame.

Timing was referenced to event code output. The duration between frame onset and event code onset was stable but inconsistent. During any given stable interval, it was approximately constant with a jitter of about +/- 1 ms, but when there were disruptions to frame refreshes, and the patch did not flicker from black to white for two or more frames, the offset reset itself. Stable frame offset values were approximately 0-10 ms. This means that offsets between frame onset and event code onset are stable over periods of tens of seconds or longer, but are not known *a priori* or repeatable between stable periods. There is no Unity-controllable way in which this can be improved on, due to the imprecision of Unity’s internal estimates of frame onset.

In summary, the timing of USE is typically precise, but contains unpredictable (non-systematic) glitches, which makes it necessary to run offline scripts to achieve sub-millisecond synchronization with frame onsets. We provide post-processing scripts that implement the accurate sub-millisecond time synchronization of frame onsets to event codes and other experimental hardware (see 5.2).

### 5.4 Gaze Classification

Gaze data from participants in the example experiment were classified using the algorithm described in 4.4.3. Table 1 shows that the velocities, amplitudes, frequencies and durations of each classified gaze period are as in previous psychophysical studies (Andersson et al., 2016; Nyström & Holmqvist, 2010; Otero-Milan et al., 2008). Furthermore, saccades exhibit the classic *main sequence* (Figure 6a), in which velocity and amplitude are linearly related, with an inflection point occurring between 10-15° (Inchingolo & Spanio, 1985).

**Table 1.** Summary statistics (means and standard errors) for gaze periods classified as fixations, smooth pursuits, and saccades, averaged across all subjects performing the example experiment. The mean velocity of saccades and peak velocities of fixations and smooth pursuits are not shown, as these are irrelevant to their characterization and tend to be misleading.

|  | Fixation | Smooth Pursuit | Saccade |
| --- | --- | --- | --- |
| **Mean Velocity (deg/s)** | 10.1 ±0.5 SE | 16.6 ±0.7 SE | --- |
| **Peak Velocity (deg/s)** | --- | --- | 210.0 ±3.6 SE |
| **Mean Amplitude (deg)** | 0.390 ±0.009 SE | 2.5 ±0.07 SE | 5.7 ±0.1 SE |
| **Rate (/s)** | 2.6 ±0.07 SE | 1.4 ±0.05 SE | 2.4 ±0.07 SE |
| **Duration (s)** | 0.35 ±0.08 SE | 0.32 ±0.01 SE | 0.04 ±0.0006 SE |

After determining fixation targets using ShotgunRaycast (4.4.3), fixations to both target and distractor objects were used to generate a heatmap of normalized fixation locations on objects, showing that participants tended to focus on object centres, though there were also a smaller number of fixations to their peripheries (Figure 6b), as one might expect in such a task (see Supplementary Materials 1 for an explanation of how normalized fixation locations were determined). Finally, participants began to preferentially fixate targets over distractors later in the block, showing that their gaze behaviour reflected their rule-learning (Figure 2e), as has been shown in previous static studies of categorization learning (cf. Blair, Watson, Walshe, & Maj, 2009; Rehder & Hoffman, 2005; McColeman et al., 2014).

## 6 Discussion

### 6.1 Overview

The *Unified Suite for Experiments* is a complete integrative suite for the development, control, and analysis of active tasks, simplifying a number of crucial challenges facing researchers interested in more realistic stimuli and possibilities of control. Its State/Level architecture supports nested hierarchies of control that enable the generation of tasks of any degree of complexity, the USE SyncBox enables reliable timekeeping and communication among experimental devices, and the offline analysis scripts enable precise time-alignment of data files and reconstruction of gaze targets.

### 6.2 Unique features of USE

Other active experimental design suites have been published (Brookes et al, 2018; Doucet et al., 2016, Jangraw et al., 2014), as well as suites that leverage active tasks for artificial learning agents (Beattie et al., 2016, Leibo et al., 2018). These offer their own advantages, and we recommend that any researchers interested in active tasks review them carefully. There are, however, several unique features of USE that may make it particularly attractive for various purposes. These include:

- ***The State/Level architecture.*** In our experience, a specialized framework for the development of flexible experimental hierarchies greatly speeds up experimental development, and makes the structure of control more apparent.
- ***The USE SyncBox.*** A generic timekeeping and communication device is a powerful tool for neuroscience research, and enables the rapid extension of USE to any new experimental hardware without a corresponding loss of temporal precision or control.
- ***Physical frame onset detection.*** USE’s ability to determine precise frame timing light sensors over the code and signal patch, and to thereby identify which specific frames were skipped or stuck during the experiment, enables the perfect reconstruction of the experimental session without relying on the game engine’s estimates of frame onset times, which do not account for skipped and stuck frames and, in our experience can sometimes be as much as 20 ms off the actual onset.
- ***Shotgun raycasts.*** So far as we are aware, all published work using eyetracking in 3D scenes either uses simple areas of interest, which do not change with regard to object shape, or single raycasts, which do not account for eyetracker imprecision or the spatial extent of the fovea. Shotgun raycasts, on the other hand, enable precise control over the degrees of visual angle surrounding gaze points in which objects are designated as targeted, without assuming anything about the underlying object shapes.
- ***Translational capabilities.*** USE enables the rapid generation and testing of slightly different versions of the same experiment, suitable for research using different populations of humans and non-humans, as well as artificial learning agents of all types.
- ***Unity online resources***. Unity is one of the most popular game engines in the world, and as such has many resources available for users of all skill levels, including tutorials, help forums, and (paid) support from Unity employees.

These advantages make USE a powerful tool for active experimentation.

Furthermore, the modular, generic and algorithmic principles underlying USE imply that it is possible for interested researchers to take any of these features that are currently unique to USE, and incorporate them into other development protocols. We hope to encourage cross-fertilization of this kind, where the best development tools and concepts from different laboratories influence those in others. For example, the gaze classification algorithm we describe in 3.4.3 was adapted from that used by Doucet et al.’s (2016) laboratory (Corrigan *et al.*, 2017), for which we are grateful.

### 6.3 Possible extensions

USE is robust and full-featured enough to be of use to any researchers interested in active tasks. However, there are many ways in which its ease-of-use, flexibility, and power could be improved. We are currently considering several of these, including:

- Creating a generic event-driven data collection system for objects in the 3D environment that automatically records their location, rotation, scale, and other alterable properties whenever these properties change. This would be combined with a generic replayer system that could read in such data files and reconstruct each frame scene, without the necessity of coding a custom replayer for each study.
- Specific wrappers for communication with a large variety of experimental hardware and software (other eyetrackers, response devices, neural acquisition devices, etc).
- Real-time analysis of light sensor data in the USE SyncBox with enough precision to enable signal output to be time-locked to frame onsets, thereby enabling real-time feedback-loops such as those used in Brain-Machine Interface contexts.
- Sub-millisecond access to system-level monitor flip commands, as is available in static experimental design suites (Brainard 1997; Kleiner et al 2007; Peirce 2007, 2008; Pelli 1997), but not in Unity. This would enable more accurate online estimation of frame onset times, and thus greatly improve frame-locked temporal synchronization in cases when using light sensors is not feasible, as is the case for, e.g., most stereoscopic goggles.
- Sub-millisecond access to inputs from serial, UDP, and TCP/IP ports, which would enable much more precisely-timed communication to/from external devices, without requiring them to be connected to the USE SyncBox. (This would still include the unavoidable delays associated with these ports, but their precision and accuracy would be much higher than in the current suite.)

USE is under active development, and we intend to update the files available for download as components are modified or created.

### 6.4 Concluding remarks

Interest in naturalistic, complex and active tasks is burgeoning, and will continue to do so. We very much look forward to seeing the effects that these tasks have on theory and method in the neurosciences, cognitive sciences, and in artificial intelligence research. We hope that the set of software and hardware components we have produced will play a small part in this process by facilitating temporally precise, well-controlled experiments in more complex and naturalistic settings.

## Supporting information

Supplemental Materials 1 - Gaze Classification

Supplemental Materials 2 - IO Timing

Supplemental Materials 3 - Normalizing Fixation Positions on Quaddle Objects

