## Supplemental Materials 1 - Gaze Classification for "USE: An integrative suite for temporally-precise psychophysical experiments in virtual environments for human, nonhuman, and artificially intelligent agents"

### Gaze classification overview

During viewing of dynamic scenes, a number of characteristic gaze behaviours can be observed, including saccades, post-saccadic oscillations, fixations, and smooth pursuits. Gaze was classified using an adaptation of a recently-developed algorithm that performs better than existing alternatives for data collected from humans viewing dynamic stimuli (Andersson, Larsson, Holmqvist, Stridh, & Nyström, 2016), and has been successfully used to classify data collected from nonhuman primates viewing both dynamic and static stimuli (Corrigan, Gulli, Doucet, & Martinez-Trujillo, 2017). Base code was provided by Corrigan et al. (2017) and modified to account for different sampling frequencies and noisier data. Details of the algorithm including our modifications are given below, and descriptions of the original algorithm are available in previous publications (Corrigan et al., 2017; Larsson et al., a2013, 2014). In brief, it classifies gaze into one of five categories: (1) saccades, (2) post-saccadic oscillations (PSOs), (3) fixations, (4) smooth pursuits, and (5) undefined, first identifying saccades and post-saccadic oscillations, and then dividing the remaining data between fixations and smooth pursuits.

### Eyetracking

Gaze was collected with a Tobii TX300 eyetracker/monitor unit, with the monitor placed above the tracker in its standard physical configuration. The active display area was 50.8 x 20.4 cm, with a resolution of 1920x1080 pixels and a 60Hz refresh rate. The eyetracker collected data at 300Hz. Participants were head free, seated approximately 60 cm from the monitor. All participants were given a 9-point calibration at the start of the experiment, which was repeated as necessary due to drift or other loss of accuracy.

### Preprocessing

Artifacts including blinks, off-screen gaze locations, one- or two-sample spikes, and any samples reported as invalid by the eyetracker were excised from the raw gaze data for each eye individually (Larsson, Nyström, & Stridh, 2013). Periods where at least 3 of 12 samples (10 of 40 ms) were invalid in both eyes were excised as well, as were any cases where the estimated gaze location was highly divergent between the two eyes (>10% of screen width, approximately 8° visual angle), along with 6 samples (20 ms) on either side of the problematic period. This divergence was mainly observed after detected blinks, and its duration usually ranged from 3-5 samples, in line with the manufacturer’s reported time to re-establish a signal after it has been lost. Location estimates were then averaged between the two eyes.

Any remaining missing data of at most 2 samples were imputed through shape-preserving cubic spline interpolation (Matlab command: interp1, argument “pchip”), and the resulting averaged position estimates were filtered using a second order Savitsky-Golay filter of length 6 (20 ms), separately for x- and y-positions (Nyström & Holmqvist, 2010). The filter was constructed with the Matlab function “sgolay”, and filtration was done through convolution with the signal. Using the sgolay function, we also obtained the first derivative filters. This was applied to the x- and y- data separately to obtain a smoothed estimate of the velocity.

### Gaze classification

The algorithm operates in multiple stages. In the first stage, it detects approximate saccadic intervals and then determines more exact saccade onsets and offsets. In a second stage, it detects PSOs. In a final stage, it classifies the remaining intervals into fixations or smooth pursuits.

To determine the approximate saccadic intervals, we note that saccades, unlike fixations or smooth pursuits, are defined by sharp increases in acceleration. Thus, we sought to identify such peaks. First, we calculated the angular acceleration by computing the angular velocity, and then smoothing this with a smoothing differentiation filter (Engbert and Kliegl 2003; Engbert and Mergenthaler 2006) of the form:

$$h\left( t \right)=\frac{f_{s}}{6}*\left[ -1-1 0 1 1 \right]$$

Next, we identified an adaptive acceleration threshold on each trial, building on work by (Nyström and Holmqvist 2010). The threshold θ is set to an initial, large value. Next, the mean and standard deviation is computed. A new threshold θ is defined as the mean + λ * standard deviation, where λ=6. Then, all points above the threshold are discarded, and the threshold is re-computed. This proceeds until the difference between successive thresholds is below a user-defined error.

However, because the algorithm as proposed uses the mean and standard deviation, it is highly biased by outliers, which in this case, are the sharp accelerations induced by saccades (Voloh et al, 2018, submitted). Thus, in the extreme case, on short trials with many saccades, the algorithm as originally defined by Nystrom et al would exit prematurely due to a large number of outliers. Reducing the lambda parameter could help alleviate this, but the optimal lambda remains a function of the number of outliers (i.e. saccades) in the data.

With this in mind, we opted to use the median and median absolute deviation as a robust estimate of the standard deviation (Voloh et al, 2018, submitted) (Leys et al. 2013), defined as:

$$\sigma_{est}=1.4826*median\left( \left| x_{i}-median\left( x \right) \right| \right)$$

Finally, putative saccadic onsets were defined as the sample of maximum acceleration in a segment that crossed the threshold. Likewise, offsets were the sample of minimum acceleration that crossed the (negative) threshold. Onsets and offsets were then matched; for each saccadic onset, we searched for the nearest occurring offset, which defined an approximate saccadic interval. Saccades less than 10ms were ignored, and saccades separated by a period of less than 20 ms were combined (Larsson et al. 2013).

Precise saccade onsets and offsets were determined for each putative saccadic interval based on two criteria, namely, (1) deviation from the main direction and (2) inconsistent sample-to-sample change in direction (see (Larsson et al. 2013; Corrigan et al. 2018)). The saccade onset was defined as the first point nearest the peak velocity for which either criteria (1) or (2) was true, looking backwards in time. The same was true for saccade offset, but looking forward in time. PSOs were estimated after each saccade in the same manner as (Corrigan et al. 2018; see also Larsson et al. 2013).

Remaining inter-saccadic intervals were then classified into either fixations or smooth pursuits. These intervals were subdivided into shorter segments by looking for periods where the sample-to-sample directions were either highly inconsistent, or highly consistent, evaluated with a sliding window Rayleigh test, as proposed by (Larsson et al. 2013). Each segment is then classified as a smooth pursuit if all 4 of the following criteria were met: (1) sample dispersion (ratio of the 1^st^ and 2^nd^ principle components) < 0.45, (2) consistent direction (ratio of the displacement relative to the fist principle component) > 0.5 , (3) total path displacement (ratio of the path displacement relative to the sum of the total trajectory) > 0.3, and (4) the magnitude of the spatial range > 1.5°. If none of the criteria were met, the segment was classified as a fixation, and when all 4 parameters do not agree, a more detailed comparison is performed (see (Larsson et al. 2013; Corrigan et al. 2018) for details).
