## Supplemental Materials 2 - IO Timing for "USE: An integrative suite for temporally-precise psychophysical experiments in virtual environments for human, nonhuman, and artificially intelligent agents"

This document describes the tests performed to characterize USE system latencies relating to the USE I/O Box. Test methods and results are summarized.

### 1. Use I/O Box Overview

The USE I/O Box is intended to perform real-time data logging as part of behavioral neuroscience experiments. A typical experiment setup is shown in Figure S1A, and the signals generated and received by the USE I/O Box are shown in Figure S1B (See also Figure 4 in the main text).

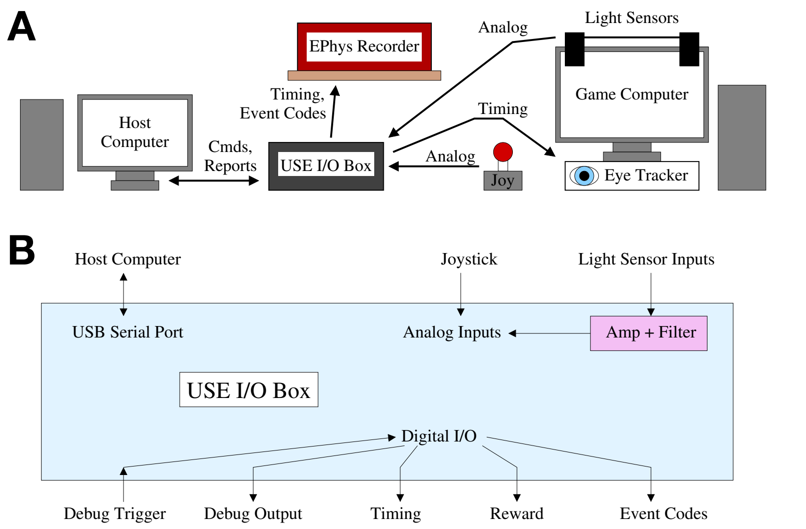

Figure S1. A) Typical experiment setup with the USE I/O box. B) Signals generated and received by the USE I/O box. See also Fig. 4 of the main text.

The USE I/O Box is implemented using an Arduino microcontroller board with a custom peripheral board attached and with custom firmware. The firmware was written with hard-real-time performance in mind; a block diagram of its implementation is shown in Figure S2.

### 2. Digital Events

The USE I/O Box enables digital signals to be generated with low latency and high precision (on the order of 0.1 ms for both). Precision and repeatability were tested by measuring timing pulses generated by the USE I/O Box; a representative oscilloscope trace is shown in Figure S3. Rise time and fall time were very much faster than the measurement scale of 0.01 ms. Jitter in the time between pulses was approximately ±0.02 ms. Signal transitions remained centered on the nominal clock interval, indicating that jitter in scheduling is much smaller than this; variation is primarily in the time between when the USE I/O Box intends to send a pulse and when it actually sends the pulse, with the intended times remaining accurately scheduled (to the limits of the measurement scale of 0.01 ms).

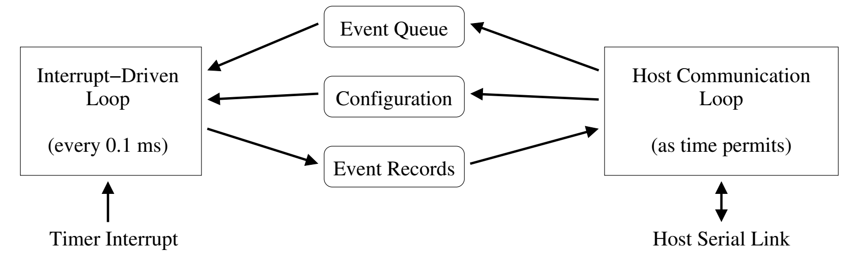

Figure S2. USE I/O box firmware block diagram

Tests were also performed of the USE I/O Box's ability to generate a digital signal when commanded via an external digital trigger; a representative oscilloscope trace is shown in Figure S3B, with the input signal below and the output signal above. Rise time was again much faster than the measurement scale of 0.01 ms. Delay between trigger and output was 0.01 ms to 0.11 ms; this was dominated by sweeping (delay progressing from one extreme to the other over about 1 s), rather than by jitter, with jitter being unmeasurably small by comparison. The magnitude of the delay is consistent with the digital sampling and event scheduling interval used by the USE I/O Box (0.1 ms); the minimum observed delay of about 0.01 ms reflects the true input delay. The sweeping observed was due to a mismatch in the clock rates of the trigger generator (another USE I/O Box) and the USE I/O Box being tested. A sweep of 0.1 ms over 1 s indicates a clock mismatch of 1 part in 10000, consistent with the clock crystals' rated accuracy of 100 parts per million.

In summary, the USE I/O Box is capable of responding to digital events within approximately 0.01 ms, with an additional delay of up to 0.1 ms due to its internal scheduling interval. It is capable of generating digital signals with scheduling precision of 0.01 ms or better, and individual event jitter of ±0.02 ms. The accuracy of the USE I/O Box's internal clock – and so the accuracy of any given measured time interval – is consistent with its rated specification of 100 parts per million.

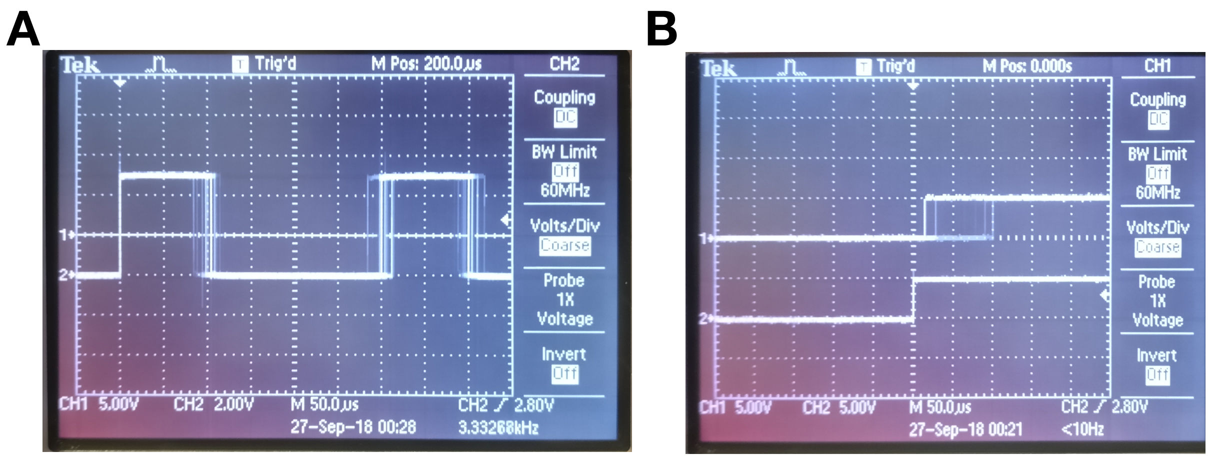

Figure S3. A) Timing signal generated by the USE I/O Box. B) Input digital trigger (below) and output pulses (above).

### 3. Analog Sampling

The USE I/O Box samples analog signals at 1 ksps. This is implemented via round-robin sampling of 5 inputs at 5 ksps, resulting in sampling time skew between nominally-simultaneous analog samples read from different inputs. Joystick inputs are directly sampled, while light sensor inputs have an analog pre-amplifier circuit. For both types of analog inputs the user can specify optional digital noise removal.

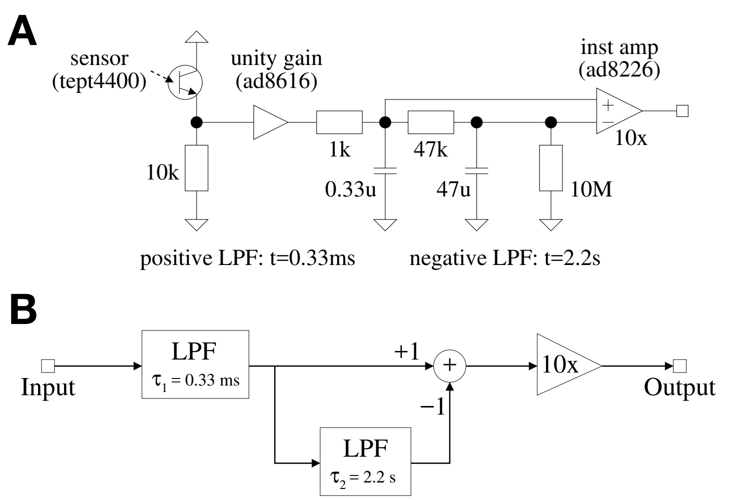


Figure S4. A) Light sensor and pre-amplifier schematic. B) Light sensor pre-amplifier block diagram.

#### 3.1 Light Filter Pre-Amplifier

The light sensor pre-amplifier performs buffering, low-pass filtering, DC subtraction, and amplification per the circuit shown in Figure S4A and the signal processing block diagram shown in Figure S4B. “LPF” blocks are first-order low-pass filters with time constants of 0.33 ms (for noise filtering) and 2.2 s (for estimating the DC level to subtract). Signals that change more quickly

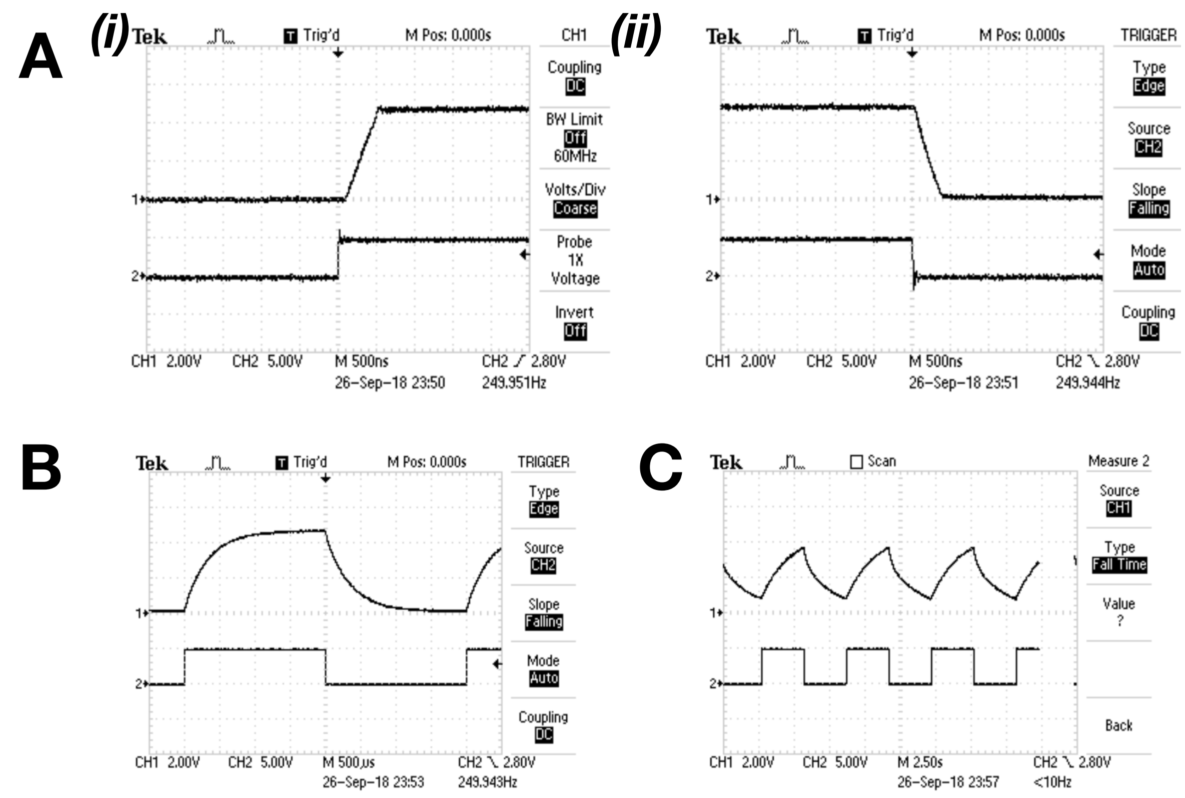


Figure S5. A i+ii) Light sensor pre-amplifier buffer output. B) Light sensor pre-amplifier fast filter output. C) Light sensor pre-amplifier slow filter output.

than these time constants turn into exponential decays with those time constants. Signals that change more slowly – with frequencies substantially below 480 Hz and 0.07 Hz, respectively – are instead delayed by their respective time constants.

Tests were performed to verify that the USE I/O Box's pre-amplifier's true output matched its predicted and intended behavior. This was done by connecting a square wave clock signal into the circuit node normally connected to the photodiode, and measuring output from the buffer amplifier, the 0.33 ms low- pass filter, and the 2.2 s low-pass filter. Representative traces can be found in Figures S5A-C. The buffer amplifier output has a rise time and fall time of about 0.5 µs; it does not introduce significant delay to signals. The filtering and DC removal low-pass filters have exponential decay responses consistent with their designed time constants of 0.33 ms and 2.2 s.

In summary, the light filter pre-amplifier is expected to delay light sensor signals by about 0.33 ms, and to take several seconds to settle after large changes to the DC light level.

### 3.2 Digital Noise Removal Filter

The digital noise removal filter is implemented using the algorithm shown in Figure S6A. Two functions are performed: low-pass filtering with a fast time constant ("fuzz removal"), and rejection of "spikes" (brief excursions with variance much greater than the average signal variance). These are illustrated in Figure S6B. The low-pass filters shown in Figure S6A are first-order exponential filters with the configuration shown in Figure S6C.

Tests were performed to verify that the digital noise filter's true output matched its predicted and intended behavior. This was done by connecting a square wave clock signal into the circuit node normally connected to the photodiode, as in Section 3.1. The digital spike removal filter was disabled, and the digital noise removal filter's time constant was set to values ranging from $2^2$ to

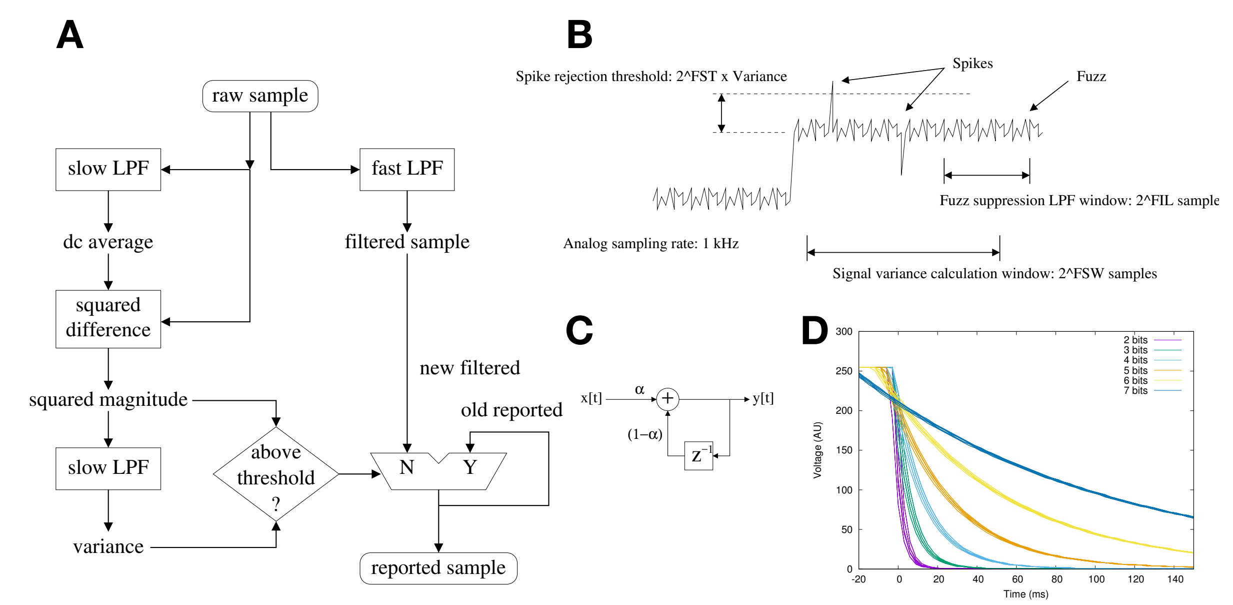


Figure S6. A) Digital noise removal filter block diagram. B) Illustration of noise features suppressed by the digital filter. C) First-order digital low-pass filter. D) Sensor response vs time for digitally filtered input.

$2^7$ samples (4 ms to 128 ms). Time skew due to the analog pre-amplifier on the light sensor input (0.33 ms) was smaller than the sampling interval and very much shorter than the digital filter time constant and so was ignored during these measurements.

Table S1. Digital Filter Time Constants

| Nominal Time Constant | Measured Time Constant |
|---|---|
| 4 ms | 3.52 ms |
| 8 ms | 7.56 ms |
| 16 ms | 15.60 ms |
| 32 ms | 31.72 ms |
| 64 ms | 63.94 ms |
| 128 ms | 127.94 ms |

Digital filtering measurement data is shown in Figure S6D. Exponential curve fits to the light sensor data were performed; the resulting nominal and actual time constants are shown in Table S1. The measured time constants are in excellent agreement with the nominal time constants.

In summary, the digital noise filter is expected to delay light sensor signals by about 4 ms if enabled with default settings ($2^2$ samples). The digital spike removal filter was not tested but uses the same architecture in its own filters; it is expected to take about 0.1 s to 0.2 s to recover from large changes to the DC light level with its default settings (a time constant of $\tau = 2^6$ samples, with a settling time of several times $\tau$).

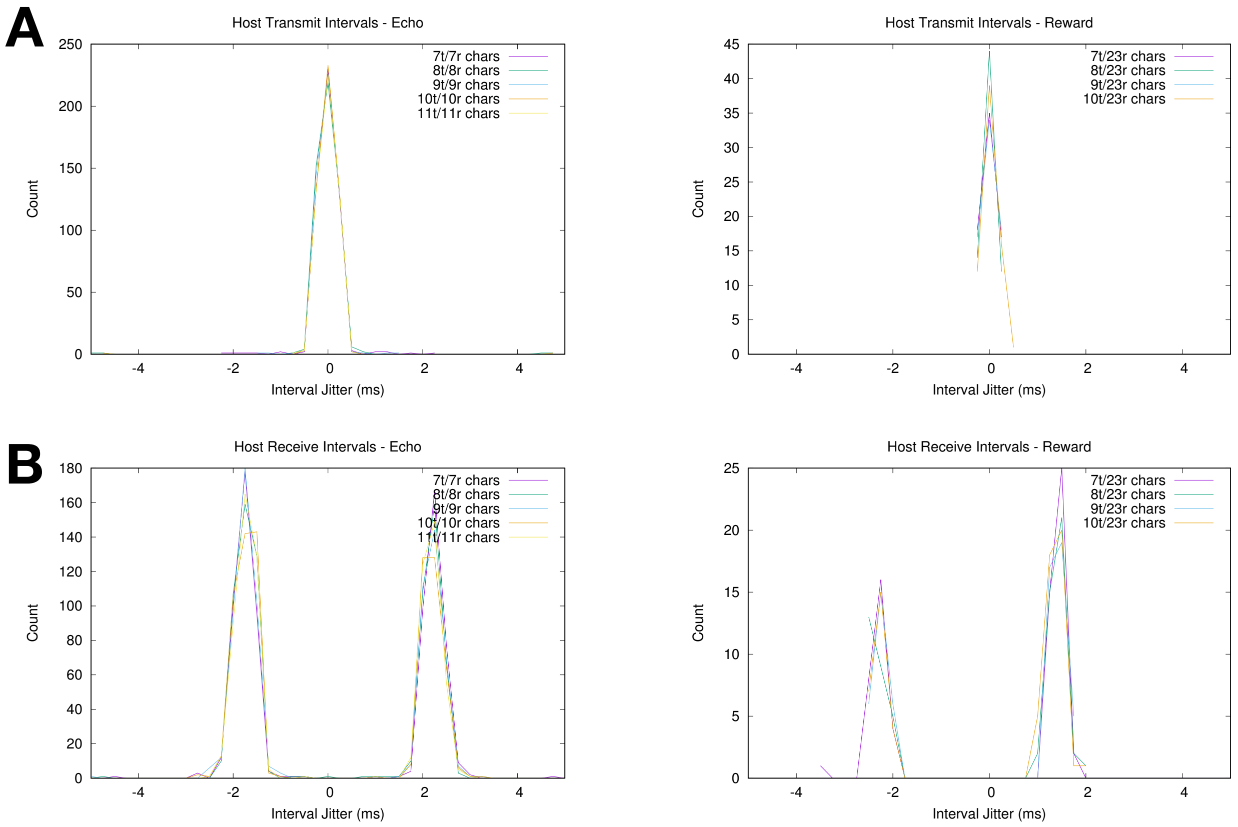


Figure S7. A) Jitter in nominal event transmission time. B) Jitter in nominal response reception time.

### 4. USB Host Link

The USB link to the host has non-negligible latency and jitter. These were measured via loopback tests, transmitting commands of known length that generate immediate responses of known length. Two such tests were performed: "echo" tests, sending a command with no effect that is echoed back to the host immediately, and "reward" tests, sending a command that generates a TTL pulse with a known event time. The nominal time at which the host sends the message and at which the host receives the message were known for all tests; for the "reward" tests, the USE I/O Box's event timestamp was also known, allowing comparison of relative drift of the host and USE timestamps.

The nominal interval between "echo" events was 10 ms (as scheduled by the host). The nominal interval between "reward" events was 150 ms (as scheduled by the host). Jitter in these intervals (deviation from the expected time) provides useful information about the delays associated with scheduling and executing communication events. Jitter in the time both types of event were nominally transmitted is plotted in Figure S7A, and jitter in the time the responses to both types of event were nominally received is plotted in Figure S7B.

Transmit time is unimodal with fitted standard deviation of 0.2 ms, and receive time is bimodal with deviation comparable to the transmit time but with displacement of ±2 ms from the mean. The smaller number – variation with a standard deviation of 0.2 ms – corresponds to the operating system's scheduler jitter (the precision with which any type of event may be queued within the host computer). The larger number – a bimodal delay separation of 4 ms – corresponds to the granularity of transmission or reception events within the operating system's USB serial driver. One or both operations may be delayed by approximately 4 ms some of the time.

Round-trip time is the time between generation of an event and reception of its response. Average round-trip times, as measured by the host, are plotted for "echo" and "reward" events in Figure S8A. Variation in the round-trip times is plotted in Figure S8B,C.

"Echo" round trip time as a function of total message size (the sum of the event message sent and the response message received) was fit to a line. This gave a fixed round-trip delay of about 2.5 ms plus a bandwidth-limited data rate of 12.1 kcps (thousands of characters per second). "Reward" data was not fit, due to excessive measurement variation (fewer trials due to the much longer event interval, and less variation in message size due to the larger fixed-length portion of the message). These numbers are consistent with the nominal channel data rate of 11.5 kcps and driver-related latencies of several milliseconds. Per-trial round-trip times vary with a top-hat distribution over a span of ±2 ms. This is consistent with the nominal transmit time varying freely but the nominal

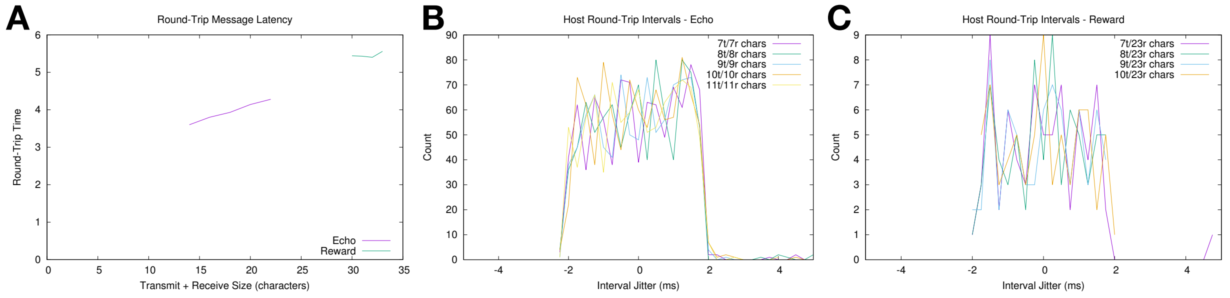


Figure S8. A) Round-trip event times as a function of total message length. B,C) Jitter in round-trip time.

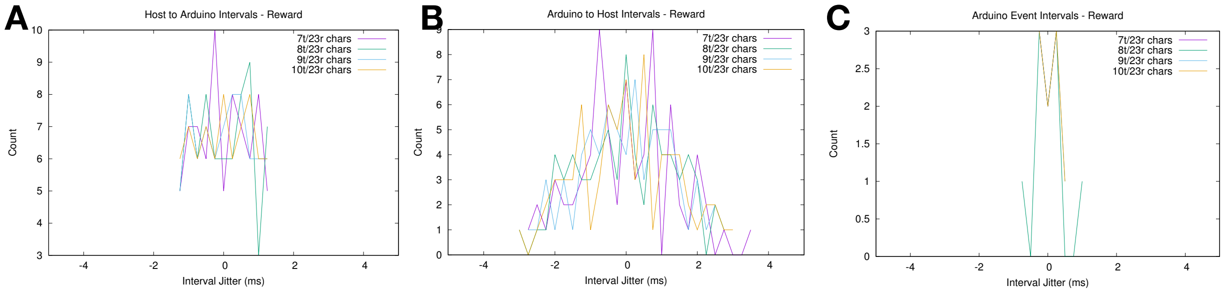


Figure S9. A,B) Host-to-USE and USE-to-host latency variation. C) Variation in USE "reward" event intervals.

receive time occurring at fixed intervals of approximately 4 ms (the time at which the USB driver reports the presence of data).

The one-way latencies between the host computer and the USE I/O Box were characterized using the "reward" tests, which provided USE timestamps in addition to host computer timestamps. The absolute one-way latencies cannot be measured directly, but variation in the differences between them are informative.

The variation in host-to-USE latencies and in USE-to-host latencies are plotted in Figure S9A and S9B, respectively. Host-to-USE latencies have a top-hat distribution with a range of approximately ±1 ms, and USE-to-host latencies have a triangle distribution with a range of approximately ±2 ms. This is broadly consistent with the "reward" test round-trip variation plotted in Figure S8B,C. Variation in the  interval between received events in the USE I/O Box is plotted in Figure S9C. This variation is small – approximately ±0.2 ms, consistent with the transmission interval variation plotted in Figure S7A. This indicates that host-to-USE latencies do not vary significantly between successive trials, but instead vary over a substantially longer timeframe.

In summary, the host operating system tested (Linux Mint 18) had event scheduling variation of approximately ±0.2 ms, which is acceptable for experiment control purposes. The operating system's USB driver, by contrast, had a scheduling granularity of about 4 ms, which introduces delays that are significant for real-time control. This is consistent with USB behavior reported by other researchers using different operating systems.
