## Supplemental Materials 3 - Normalizing Fixation Positions on Quaddle Objects for "USE: An integrative suite for temporally-precise psychophysical experiments in virtual environments for human, nonhuman, and artificially intelligent agents"

### Determining the distribution of fixations onto objects

All gaze samples collected during an experimental session were aligned with corresponding frames (see section 3.4.1 of the manuscript). The entire session was then replayed (3.4.2), and on each frame the targets of each gaze point in the scene were determined. This was done using a ShotgunRaycast (see section 3.4.3), set such that the maximum distance between the origins of rays on the camera surface was 0.2° of visual angle, and the radius of the outer circle was 1° of visual angle, using the sample-by-sample distance of participants’ eyes from the eyetracker. The proportion of rays within each gaze point’s ShotgunRaycast that hit one of the two Quaddle objects in each trial was recorded. If this proportion was greater than 0, this gaze point was understood to be targeting this Quaddle. For each fixation during the experiment (see 3.4.3 and Supplementary Materials 1 for how fixations were determined), it was deemed to be targeting any Quaddle that was being hit by multiple constituent gaze points. (This is a liberal definition of fixation targeting, which we used it due to its simplicity, and the negligible number of ambiguously-targeted fixations we observed in informal reviews of the data).

For each frame during a fixation of a Quaddle, we determined the XY-position of gaze on the screen S_fix_. We then extracted three other quantities, including the position of the player P_player_ in world units, the position of the centre of the object in world units P_object_, and the position of the centre of the object on the screen S_object_. We extracted the 2D vector D_fix_ = S_object_ - S_fix_, representing the position of the fixation on the screen relative to the object centre. However, the two-dimensional size of the silhouette of an object is inversely proportional with distance. To account for this, we scaled the magnitude of this vector by the distance of the object in the world, d_object_ = |P_player –_ P_object_|. Fixations to objects that were closer than 4 world units to the player were not included, as the relationship between distance and two-dimensional size was non-linear below this distance. Thus, for each fixation frame, we extracted one normalized vector, D_norm_ = D_fix_ / d_object_, representing the normalized screen position of the fixation relative to a fixated object.

To generate Figure 6B of the main text, we created images using a standard Quaddle presented 4 world units away from the camera, at the same angle as participants would have viewed it. We scaled the image into the same units as D_fix_, by measuring the width (from one arm end to the other) and height (from bottom to top) of the objects relative to the screen size. We binned D_norm_ into a 2D histogram with 30x30 equally spaced bins, and overlaid this heatmap over the quaddle image.
